# Selective encoding of priors for flexible categorization but not Bayesian inference in the frontal eye field

**DOI:** 10.1101/2024.12.31.630950

**Authors:** Divya Subramanian, John M. Pearson, Marc A. Sommer

## Abstract

Expectations or prior beliefs about the world have been shown to modulate sensory processing both at the behavioral and neural levels. Bayesian models predict that such priors compensate for input uncertainty to optimize sensory judgments. Although Bayesian behavior is prevalent across sensorimotor systems, the relationship between priors and Bayesian inference is not obligatory. Priors may simply shift one’s internal decision boundaries without interacting with sensory uncertainty at all. We recently showed that humans and monkeys use both Bayesian and non-Bayesian strategies when reporting judgments of visual stability across saccades, despite using priors in both cases. While they increased prior use to compensate for internal, movement-driven sensory uncertainty in a Bayesian manner, they *decreased* prior use when faced with external, visual image uncertainty. The latter, “anti-Bayesian” pattern was best explained by a model in which category boundaries were adjusted by the prior but susceptible to image noise. Here, we recorded neural activity in the frontal eye field (FEF), a prefrontal region important for visuosaccadic behavior, while toggling between subjects’ prior use for Bayesian and anti-Bayesian behavior via trial-by-trial manipulation of the two uncertainty conditions. First, we found that FEF activity signaled the priors in both conditions. The prior-related modulation of activity, however, predicted only the anti-Bayesian, categorization behavior. The results suggest that neural activity in the FEF reflects the use of a flexible decision boundary for the perception of visual stability and, more generally, that neural mechanisms for Bayesian inference and visual categorization are dissociable and distributed in the primate brain.

**Significance:** Appreciating a visual scene depends not only on retinal input, but also on priors about the world. Foreknowledge interacts with visual inputs to improve reactions and decisions. One way the brain combines priors and inputs is by using Bayes’ rule to model optimal outcomes. A simpler way is by categorizing inputs with prior-adjusted boundaries. Here, we tested how neurons in primate frontal cortex use priors: for Bayes’ rule, or for flexible categorization? A key feature of the study was to use a single perceptual task that was varied trial-by-trial to yield either Bayesian or categorization behaviors. We could then establish which behavior the neurons encoded. The implications extend beyond visuomotor behavior to broader neurocomputational mechanisms of prior use for cognition.

## Introduction

Priors about the environment are known to modulate sensory-guided behavior and its associated neural activity (Summerfield and de Lange, 2014; Bansal et al., 2018; de Lange et al., 2018). Bayesian models, which predict that priors may compensate for uncertainty in sensory inputs, have been successful at explaining many behaviors (Jacobs, 1999; Ernst and Banks, 2002; Weiss et al., 2002; Knill and Saunders, 2003; Kording and Wolpert, 2004; Girschick et al., 2011; Fetsch et al., 2012; Darlington et al., 2017; Jazayeri and Shadlen, 2010). However, prior use does not necessarily imply Bayesian inference. Priors may simply alter behavior in a flexible, context-dependent manner without compensating for uncertainty as Bayes’ rule demands (Green and Swets, 1966).

We recently demonstrated this dissociation between priors and Bayesian inference for visual perception across saccadic eye movements (Subramanian et al., 2023). The challenge of maintaining coherent perception across saccades is fundamental to primate behavior. Each saccade causes the retinal image to move, requiring the visual system to disambiguate saccade-induced retinal shifts from external object movement. One way to solve this problem is by using a copy of the saccadic command (corollary discharge) to predict the expected visual inputs after a saccade (Sommer and Wurtz, 2008). Inputs that match the prediction can be attributed to one’s own movement and mismatches to external movement. This process is susceptible to two major sources of sensory uncertainty: 1) internally generated noise caused by a degradation of visual processing around the time of saccades, called saccadic suppression (*saccade-driven noise;* Zuber and Stark, 1966), and 2) external noise in the visual input *(image noise*). We previously found that humans and monkeys use priors about the probability of external object movement (Rao et al., 2016; Subramanian et al., 2023). However, while prior use increased with internally generated uncertainty caused by their own eye movements (consistent with Bayesian predictions), prior use *decreased* with additional image noise. This “anti-Bayesian” pattern was consistent with the flexible adjustment of visual category boundaries across prior conditions.

Here we studied the neural correlates of noise-affected visual stability judgments in the frontal eye field (FEF), a prefrontal region important for visual processing across saccades. We examined whether FEF activity encodes priors and, if so, whether the neural prior signals exhibit modulation more consistent with Bayesian inference or categorization strategies (Fig. 1a). The FEF is a primary candidate region for implementing the two computational strategies for the perception of visual stability across saccades (Fig. 1b; Sommer and Wurtz, 2008; Cavanagh et al., 2010; Crapse and Sommer, 2008; Hall and Colby, 2011; Rao et al., 2016; Subramanian et al., 2019; Cavanaugh et al., 2016). Some FEF neurons receive corollary discharge signals that remap their visual receptive fields to their post-saccadic locations before the onset of the saccade (Umeno and Goldberg, 1997; Sommer and Wurtz, 2006; Wang et al., 2024; Jin et al., 2025). Since remapping allows neurons to sample the same region of space before and after a saccade, it can serve as the basis for predicting the expected post-saccadic visual input (Hall and Colby, 2011; Wurtz, 2018). Moreover, the FEF receives retinocentric information (Bruce and Goldberg, 1985) and signals the mismatch between this input and its predictions (Crapse and Sommer, 2012).

**Figure 1.**
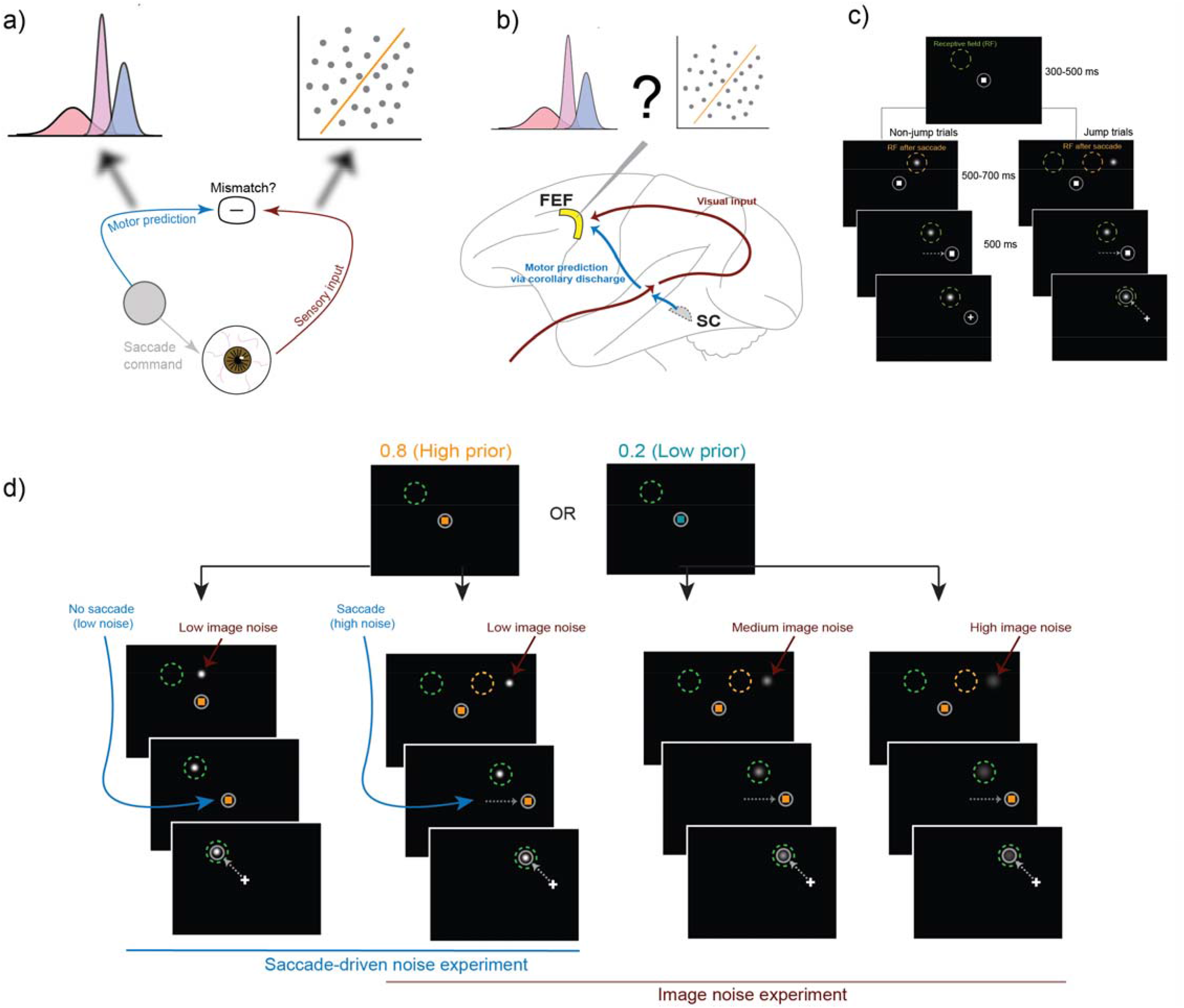
Experimental question and design. a) To judge whether a sensed retinal shift was caused by one’s own eye movements or external object movement, the visual system can use a copy of the saccadic command to predict its sensory consequence. Inputs that match this prediction can be attributed to self-movement and mismatches to external object movement. We previously showed that subjects use Bayesian inference to resolve uncertainty caused by one’s own movement (left fuzzy arrow) but flexibly shift a discriminative category boundary to account for uncertainty caused by external image noise (right fuzzy arrow). b) We recorded from single neurons in the frontal eye field (FEF), which receives visual and saccade-related inputs, to test for neural correlates of the Bayesian and discriminative strategies. SC, superior colliculus. c) Typical spatial configuration in the task. Subjects initiated a trial by fixating on a central square. White circle depicts the eye position. A Gaussian visual probe was presented at this time. On jump trials, the probe was presented near the center of the post-saccadic receptive field (orange circle) and for non-jump trials, it was presented at the center. After a brief fixation period, the square moved to the target location and subjects followed it with a saccade (dashed white arrow), during which the probe was moved on jump trials. For both jump and non-jump trials, the probe was at the center of the receptive field (green circle) when the subject made a response. A perceived jump was then reported with a second saccade to the probe. Perceived non-jumps were reported by maintaining fixation. d) Priors were indicated to the subject on every trial by the color of the fixation square. The two experiments included 4 noise conditions in total, with one condition in common between them. The saccade-driven noise experiment included two conditions, one where the subject made an intervening saccade (as shown in c) and another where they maintained central fixation during probe movement. Visual noise was manipulated by the width of the Gaussian probe. Left-most column: no-saccade condition with low saccade-driven noise. Left-middle, right-middle, and right-most columns: conditions with an intervening saccade (with saccade) but with low, medium, and high image noise respectively. The condition with a saccade and low image noise serves as a common condition between the two experiments. Target, probe, and receptive field sizes are all schematized for clarity and not drawn to scale.

We recorded FEF neurons while subjects performed the image- and saccade-noise tasks, interleaved trial-by-trial. First, we found that FEF neurons were selective for priors. Then we evaluated whether the neuronal prior selectivity increased with saccade-driven noise (the Bayesian prediction) or decreased with image noise (consistent with visual categorization). We found that FEF activity did not significantly correlate with increasing prior use in the saccade-driven noise task, but it did correlate with and predict decreasing prior use in the image noise task. The FEF’s selective participation in visual categorization but not Bayesian inference, even though both strategies were used behaviorally, suggests that they are mediated by dissociable, distributed circuits.

## Materials and Methods

### Surgical preparation and electrophysiological recording

Two rhesus macaques (Monkey S and Monkey T, the same male animals used for the studies reported in Subramanian et al., 2023) were used for procedures described in this manuscript. Animals were brought into the lab in custom-made chairs (Crist Instruments, Hagerstown, MD) and their heads were stabilized using a head-post that attached to both the chair and a surgically implanted socket (Crist Instruments, Hagerstown, MD) on the skull. The socket, a stereotaxically positioned recording chamber over the frontal eye field (FEF), and monocular scleral search coils (Robinson, 1963; Judge et al., 1980) were implanted in an aseptic surgical procedure. All surgical and experimental procedures were performed in accordance with protocols approved by the Duke Institutional Animal Care and Use Committee. They were positioned 60□cm from an LCD monitor (1920□× 1080, 144□Hz) on which visual stimuli were presented.

Neurons in FEF were recorded by placing a 1 mm x 1 mm grid within the recording chamber and advancing sterilized tungsten microelectrodes with impedances ranging from 750-950 kΩ (FHC, Bowdoin, ME) through a guide tube. The FEF was identified anatomically in each animal using structural MRI and by the visual and presaccadic response properties of recorded neurons. Neurons were extracellularly isolated by identifying the auditory and visual signatures of individual waveforms using an audio speaker and oscilloscope respectively. Continuous data for all recorded signals (neural signals sampled at 10,000 Hz and eye position sampled at 1000 Hz) were stored. Neural activity was passed through a 60 Hz notch filter. Online, the neural activity of one or more putative single neurons was recorded, aligned to behavioral trial events, and monitored using rasters and spike density functions using Spike2 (Cambridge Electronic Design, Cambridge, UK) and custom Matlab code (Mathworks). This online monitoring aided in ensuring sustained isolation of specific action potential waveforms throughout the session, which could last 1-3 hours. Offline, action potentials were sorted to confirm single neuron origins using a Principal Components Analysis based clustering algorithm in Spike2. All other analyses were performed in Matlab (Mathworks).

### Receptive field mapping

After a neuron was isolated in each session, we first mapped its visual, motor, or visuomotor spatial receptive field in two delayed saccade tasks. On every trial, the animal first fixated a white, central square (1° x 1°) for a randomized duration of 300-500m. All randomized delays reported in this manuscript were drawn from uniform distributions. A target square (also white and 1° x 1°) was then presented concurrently with the fixation square for a delay of 500-700ms. After the delay, the fixation square was extinguished cuing the animal to make a saccade to the target. Upon target acquisition, the animal was further required to fixate on the target for an additional 500 ms after which, fluid reward was delivered. Since the first occurrence of the target, i.e., its onset, is separated by 500-700ms from the time that the animal makes a saccade to it, we can distinguish the neuron’s visual response (typically 50-100ms after target onset) from its motor response (typically 150-100ms before the saccade). We first mapped the neuron’s responsiveness to the target placed in one of 8 potential directions: 4 cardinal directions and 4 along the diagonal. The targets were placed at a retinal eccentricity that was most likely to match neurons’ preferences in the subregion of FEF. Typically, this was at 10°. Next, based on the neuron’s direction preference, we then ran the amplitude task, where the target was restricted to the preferred direction but presented at one of four amplitudes: 2°, 5°, 10°, and 20°. The putative center of the receptive field was inferred based its response properties in the direction and amplitude tasks and used to guide the placement of the probe in the Saccadic Suppression of Displacement (SSD) task.

#### Experimental design and statistical analyses

##### Saccadic Suppression of Displacement (SSD) task

The overall goal of the current manuscript was to record the activity of FEF neurons while we tested the use of learned priors to compensate for internal, saccade-driven or external, image noise in the context of a saccadic suppression of displacement task. Animals made saccades and reported whether a visual stimulus in the environment moved during the saccade.

On each trial, a fixation square (1ºx1º) first appeared at the center of the screen. After fixation had been acquired and maintained for a randomized duration of 300-500ms, the visual probe appeared at one of the 4 locations on the screen for 500-700ms. The monkey was required to maintain fixation on the central fixation square for that duration, after which the fixation square was replaced by the saccade target (1ºx1º) indicating to the animal they could make a saccade. Saccade initiation (defined as the time the eye crossed a threshold set at 20% of the saccade length) triggered the displacement of the probe on some trials. Animals were further required to maintain postsaccadic fixation for 500ms after which the saccade target was replaced by a white cross in the same location. To report that the probe had moved during the saccade, the monkey was required to make a saccade to the probe within 500ms and then fixate on it for 400ms. To report that it had remained stationary during the saccade, the monkey had to remain fixated on the cross for 1000ms. The precise timing of stimulus presentation was verified with a photodiode taped to the top left corner of the monitor, where a white square (invisible to the monkey) was flashed within the same frame as the measured stimulus.

The probe was initially placed at a location determined by the displacement and was always displaced parallel to the intervening saccade such that it landed at the center of the cell’s expected receptive field after the planned saccade. The intervening saccade vector was chosen on a session-by-session basis to be away from the receptive field. The saccade target location was jittered by 1° trial-by-trial, so that its absolute location did not correspond with a jump or non-jump response. For trials in which no saccade was made (described below), the probe landed in the center of the receptive field. Displacements were drawn from relatively broad and narrow one-dimensional Gaussian distributions in the movement (μ = 0º, σ = 2.5º) and non-movement (μ = 0º, σ = 0.2º) conditions, respectively. Positive displacements were either rightward or upward, and negative displacements were leftward or downward. For each session, four to six discretized displacements were chosen to allow for a comparison of neural responses across prior and noise conditions with variability minimized across all other parameters. The displacements were not the same across sessions.

Priors (0.2 and 0.8) were cued by the color of the fixation and target squares. For monkey S, green squares meant that the probe had a 0.2 probability of being displaced while magenta squares indicated a 0.8 probability of displacement. For monkey T, blue squares were associated with a 0.2 probability of displacement while orange squares were associated with a 0.8 probability of displacement. Animals were trained on priors over multiple sessions using performance-based feedback. External, image noise was manipulated by the width of the Gaussian probe, which had one of three possible standard deviations: 0.5º (“low noise”), 1.25º (“medium noise”), and 2º (“high noise”) for Monkey S and 0.5º (“low noise”), 1.25º (“medium noise”), and 1.75º (“high noise”), for Monkey T. To manipulate internal, saccade-driven uncertainty, we added a condition to the experiment where monkeys did not make a saccade. The purpose was to eliminate a major form of saccade-driven sensory uncertainty, the saccadic suppression of visual signals. The monkeys remained fixated in the center while the Gaussian, visual probe was displaced. This no-saccade condition served as the “low noise” condition and was compared to a “high noise” condition where animals made a saccade. The Gaussian probe had low image noise (i.e., a standard deviation of 0.5º) in these conditions. The temporal structure of the no-saccade trials was identical to the trials with a saccade.

The low image noise condition with a saccade served as the common condition between the saccade-driven and image noise experiments. Thus, there were a total of four noise conditions at two prior levels. Conditions with low image noise also included a 0.5 prior condition with a white fixation and target square (data not shown but used for model fitting). All trial types were equally likely and randomly interleaved.

##### Data inclusion

We recorded from 91 FEF neurons (55 from Monkey S across 40 sessions, and 36 from Monkey T across 24 sessions). For each session, a small minority of trials in which the probe did not land in its final position before the end of the intervening saccade were excluded from analysis. As in Subramanian et al. (2023)^26^, we measured the timing of the probe landing using the response of a photodiode to a white square, invisible to the monkey, presented in the top left corner of the screen simultaneously as the probe. We verified the maximum duration of a frame as being 7ms from top left to bottom right using a second photodiode. Since the probe was presented at various locations on the screen, we set the most conservative criterion such that the photodiode timestamp had to be at least 7ms before the detected end of the saccade. We limited our analyses to neurons recorded on behavioral sessions in which the animal completed at least 10 valid trials in each condition of the experiment. This resulted in a total of 79 included neurons (53 from Monkey S across 39 sessions and 26 from Monkey T across 17 sessions). For control analyses performed only on trials where the displacement was 0, we used neurons for which the animal completed at least 5 valid trials with displacement = 0 in all conditions. This resulted in the inclusion of 64 total neurons.

##### Behavioral prior use and comparison to model predictions

We quantified animals’ “prior use” as being the difference in the proportion of trials in which they reported “jumped” across prior conditions. We compared the pattern of their prior use data across noise conditions to the simulated predictions from two models evaluated in in Subramanian et al. (2023): a Bayesian ideal observer model and a model combining the outputs of a Bayesian model and a perceptron-based category-learning model.

###### Bayesian ideal observer model

The ideal observer makes a probabilistic decision about binary variable, *J*, indicating whether the target jumped or not. ¬*J* indicates that the target did not jump. Since the true displacement is experimentally drawn but not available to the observer, they make this decision given the *perceived* displacement, 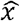. The decision is based on the relative probabilities of the target having jumped or not jumped given the perceived displacement,

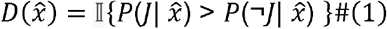

where 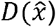 is the decision given the perceived displacement, 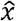, and is determined by a binary indicator function, 𝕀. 𝕀 = 0 (no jump) if the condition in braces is not met. Otherwise, 𝕀 = 1 (jumped); 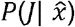 is the probability that the probe jumped given 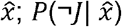 is the probability that the probe did not jump given 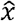.

Using Bayes’ rule for the condition within braces in Eq. 1,

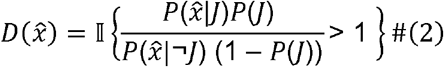

The simulated decision of the ideal observer, however, must be compared to the subject’s responses. We do not have access to the subject’s perceived displacement, but instead can only infer their decision given the true experimental displacement, *x*. We assume that the perceived displacement is a Gaussian random variable where the mean is the true displacement, and its variance given by the width of the blob on that trial:

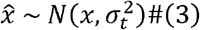

where 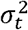 is the variance of the target.

The decision given the true displacement can thus be modeled as,

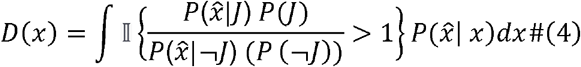

That is, the decision value given the true displacement, *D*(*x*), is the integral of the perceived displacement distribution that falls above the point at which the indicator function, 𝕀, is non-zero.

Based on the distributions used in the experiment,

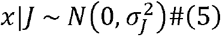

and

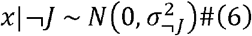

Since 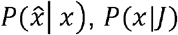, and *P*(*x*|¬*J*) are Gaussian distributions, we integrate over *x* such that,

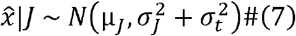

and

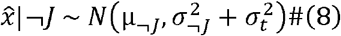

Thus, the expression inside the indicator function in Eq. 4, when replaced with the appropriate Gaussian probability density functions, equals,

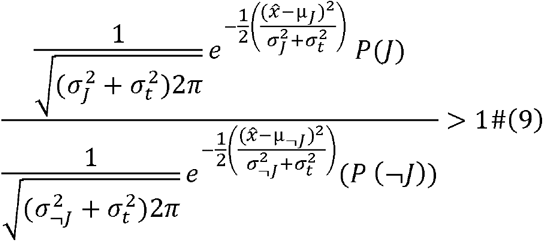

Taking the log on both sides provides the condition under which the indicator function is > 0,

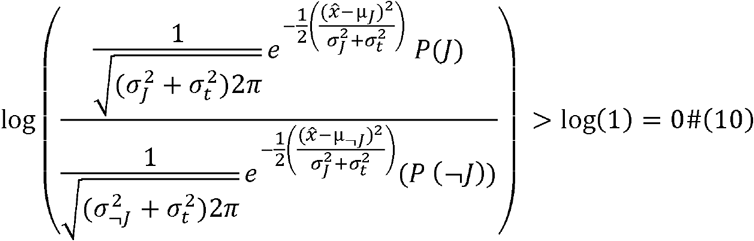

That is,

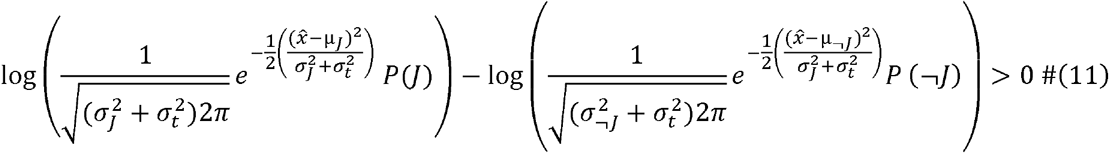

Rearranging terms, the indicator function is > 0 when

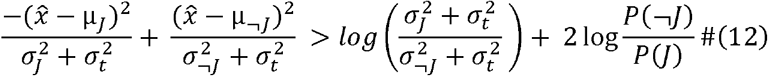

When μ_*J*_ = μ_¬*J*_ = 0 as in our experiment, the indicator function is > 0 when 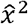 is greater than a criterion value, 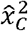, defined as,

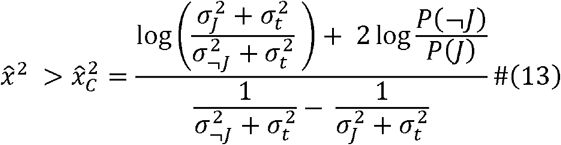

Since 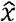 is a Gaussian random variable, 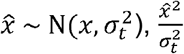 is a non-central chi-square random variable, 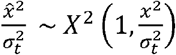. Thus, the decision, *D*(*x*), can be modeled as the integral of a non-central chi-square distribution that lies above the criterion, 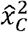. That is,

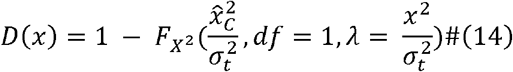

where 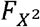 is the cumulative distribution function of *X*^2^ with degrees of freedom, *df* = 1 and gamma, 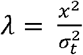 up to 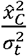.

Based on the results in Subramanian et al. (2023), we simulated the three levels of image noise by varying the values of *σ*_*t*._ and set them at 0.4º, 0.65º, and 0.9º for low, medium, and high noise respectively. Saccade-driven noise was simulated by varying the width of the non-jump distribution, *σ*_¬*J*_ We set *σ*_¬*J*_ = 0.1º for the no-saccade condition, and *σ*_¬*J*_ = 0.75º for the conditions with an intervening saccade. The width of the “jump” distribution, *σ*_*J*_, was set to 5º for all conditions. Simulation parameters for both models were chosen to best illustrate the predicted pattern of prior use across the four noise levels, and simulated results are expressed in arbitrary units.

###### Category-learning model

We set up a two-layer neural network to classify displacements into categories of “jumped” and “did not jump” and simulated its performance under experimental conditions. The input layer consisted of units representing continuous displacements and the output layer had two units: “jump (J)” and “no jump (NJ).” For ease of computation, continuous input displacements were discretized into bins of 0.1°. Displacements ranged from 0 - 7.5°. That is, there were 75 input units for each network. Sensory noise was simulated as a Gaussian distribution of input unit activation, truncated at the two ends of the input range (0 and 7.5), such that the total activation of input units was always 1. On each trial, the distribution was centered on the true displacement for the trial and the width of activation was determined by the sensory noise level. Each input unit was connected to both output units. The activation of each output unit was the *weighted* sum of inputs, i.e.,

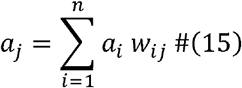

where *a*_*j*_ is the activation of the output unit, j; *a*_*i*_ is the activation of the input unit, i; and *w*_*ij*_ is the weight of the connection between input unit, i, and output unit, j. The final output on each trial was the normalized activation of the “jump” and “no jump” output units such that the output for each unit was bounded between 0 and 1:

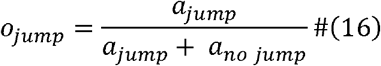

and

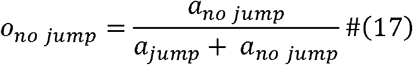

where *o*_*jump*_ is the final output of the “jump” unit, *o*_*no jump*_ is the final output of “no jump” unit, and *a*_*jump*_ and *a*_*no jump*_ are activations of the “jump” and “no jump” output units, respectively.

There were separate sets of inputs for each prior. During the training phase, weights between inputs and outputs were updated using a perceptron-like, error-based learning rule:

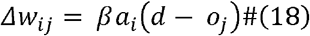

where *Δw*_*ij*_ is the change in weights between input unit, i, and output unit, j; *β* is the learning rate, *a*_*i*_ is the activation of the input unit, i; *o*_*j*_ is the final output of unit j, and d is the desired state of output unit, j. d was set to 1 for the jump output and 0 for the non-jump output on trials where the probe jumped, and vice versa for non-jump trials.

###### Combined model

The final output of the combined Bayesian and perceptron-like category-learning model was obtained using a weighted combination of the output of each model individually:

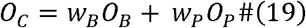

where *O*_*C*_ is the output of the combined model, *O*_*B*_ is the output of the Bayesian model, *O*_*P*_ is the output of the perceptron model, and *w*_*B*_ and *w*_*P*_ are the weights assigned to the Bayesian and perceptron models, respectively. Further, the weights of the two component models added up to 1:

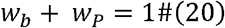

We simulated behavior in the saccade-driven and image noise experiments by modeling saccadic noise within the Bayesian ideal observer model and image noise in the perceptron model as described in Subramanian et al. (2023). Each simulation consisted of 2000 trials. The width of the “jump” distribution, *σ*_*J*_, was set to 2.5º. We set *σ*_¬*J*_ = 0.5º for the no-saccade condition, and *σ*_¬*J*_ = 1º for the conditions with an intervening saccade. Image noise (*σ*_*t*_) was set to a constant 0.1º for all conditions in the Bayesian model. For the perceptron model, low, medium, and high image noise were simulated at 0.1º, 1º, and 2º respectively. For the first 1000 trials, the model was trained under conditions similar to those experienced by the subjects. The weights between the inputs and outputs were trained in the low image noise, with saccade condition using a learning rate of 0.5. The remaining 1000 trials formed the test phase with the 4 noise conditions described in this manuscript. For these trials, the weights remained constant, and saccade-driven and image noise were simulated as described above. *w*_*b*_ and *w*_*P*_ were set to 0.1 and 0.9, respectively. We chose these values of *w*_*b*_ and *w*_*P*_ for simulations to qualitatively best match the observed data shown in Fig. 2g. The predicted outputs of the model at *w*_*b*_ = 0.25, *w*_*b*_ = 0.5, *w*_*b*_ = 0.75, and *w*_*b*_ = 0.9 are additionally shown for comparison in Figure S2.

**Figure 2.**
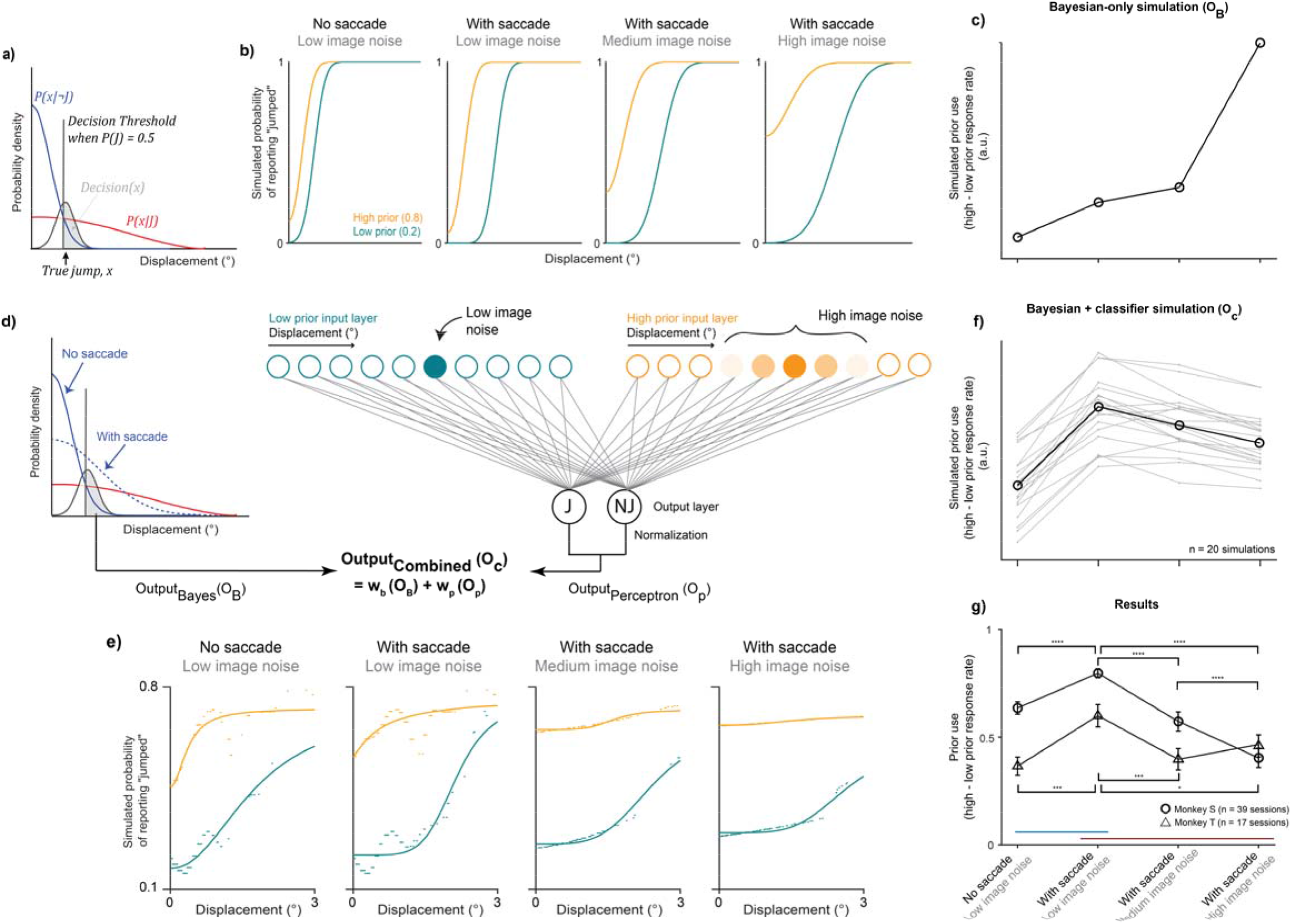
Behavioral models and results. a) Bayesian ideal observer model schematic. Overlapping blue and red distributions depict the Gaussian non-jump and jump distributions, respectively, both centered on displacement = 0. High saccade-driven noise analogous to saccadic suppression is modeled by increasing the width of the non-jump distribution. The black distribution depicts the Gaussian distribution associated with probe. The probe distribution is centered on the true displacement (black arrow) with its width determined by the amount of image noise. Black vertical line in the left-most panel depicts the decision threshold, or the point at which a given displacement was likely drawn from the non-jump vs. jump distributions, when the relative probabilities of being drawn from either are equal (i.e., the prior = 0.5). The value of the model output, i.e., the “decision”, for each displacement and prior condition is determined by the integral of the probe distribution over displacement values greater than the decision threshold (gray shaded area in left-most panel). Prior and uncertainty simulations are not schematized here but described in detail in Subramanian et al., 2023. When the prior is less than 0.5, the threshold moves to the right and when greater than 0.5, to the left. Saccade-driven noise was simulated by a widening of the non-jump distribution, and image noise by a widening of the target distribution. b) The Bayesian model predicts that subjects’ probability of responding “jumped” increases with a high prior relative to a low prior (orange vs. blue lines), and that this difference increases steadily with increasing saccade and image noise. c) Prior use, quantified as the difference in response rates between the high and low priors, across the four noise conditions (see x-axis labels in panel g; O_B_, output of the Bayes model). d) Combined Bayesian and perceptron model schematic. Model schematic. Left hand side depicts the Bayesian observer model from Fig. 2a. High saccade-driven noise, i.e., in conditions with an intervening saccade, is modeled by increasing the standard deviation of the non-jump distribution relative to the jump distribution. The image noise level in the Bayesian model was held constant at the lowest level. Medium and high image noise were applied instead to the input layer of a two-layer perceptron-like category-learning network (right hand side). The input layer consisted of displacement units with one set each per prior. Input activation was centered on a given displacement and the width of activation was determined by the image noise level. Example low image noise activation shown on the left and high image noise on the right (intensity of the color reflects the width of the activation). The total input activation on each trial always summed to 1. Activation of the “jump” and “no jump” units in the output layer, which were determined by a weighted sum of activated inputs, were normalized for a perceptron model output. The final, combined model output was a weighted sum of Bayesian and perceptron outputs. e) Example psychometric curves across the four noise conditions from a single simulation. Curves were fit to model response rates on individual trials shown as dots. f) The combined model predicts that prior use increases with increasing saccade-driven noise but decreases with additional image noise (see x-axis labels in panel g; O_C_, output of the combined model). We quantified prior use as the mean difference in response rates between priors for displacements ranging from 0-2°, to restrict the quantification to displacements that were prevalent across both the low and high prior conditions. Thin gray lines depict each simulation. Thick black line shows their mean. g) Data from the two monkeys were consistent with the predictions of the combined Bayesian and categorization model. Error bars indicate the standard error of the mean. Prior use increased with saccade-driven noise, consistent with Bayesian predictions, but decreased with additional image noise, consistent with the use of a categorization model. Asterisks at the top for Monkey S and on the bottom for Monkey T. *, p < 0.05; ***, p < 0.001; ****, p < 0.0001.

##### Model fitting

We fit the combined model to subjects’ behavioral responses on a session-by-session basis using maximum likelihood estimation, i.e., by minimizing negative log likelihood between the model’s output and subjects’ binary responses. To simulate the learned weights between the displacement inputs and the “jump” and “non-jump” outputs for each prior condition in the perceptron component, we used the value of the psychometric curve fit to all data in that prior condition for each session.

We allowed 12 parameters to be free: 2 noise types for all 4 experimental conditions each, the width of the “jump” distribution, and 3 priors. Parameter optimization was implemented using MATLAB’s *fmincon* function.

##### Neuronal firing rate analyses

###### Normalization

To visualize population peri-stimulus time histograms aligned to specific task events, we first normalized the activity of each neuron and then averaged their firing rates. Firing rates were z-score normalized by binning spikes across all task conditions into non-overlapping 20ms bins, subtracting the mean of the bins, and dividing by their standard deviation. Normalized firing rates were additionally convolved with 20ms Gaussian kernels.

###### Non-parametric statistical significance test and effect size calculation

Firing rates in the high and low prior conditions were evaluated for statistical significance using the Wilcoxon rank sum test, and the effect size of the difference was quantified using the Wilcoxon effect size. The Wilcoxon effect size is given by 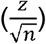, where z is the z-statistic used by the rank sum test and n is the total number of trials (low prior + high prior). A positive effect size indicated that firing rates in the high prior condition were greater than in the low prior condition, and vice versa for a negative effect size.

###### Mixed-effects regression model

To compare neuronal prior use with behavioral prior use while simultaneously accounting for all other experimental variables, we ran two mixed-effects regression models on neurons with significant prior effects in the common condition: one with the neuronal firing rates as the outcome variable and one with the behavioral responses (“jumped” or “did not jump”) in the same sessions as the outcome. The models were run on the data used to analyze neuron-behavior correlations in Figure. 5f-h and Figure S8, i.e., the neurons for which there was a significant prior effect in the reafferent epoch of the common condition (i.e., 0 – 800 ms from the landing of the intervening saccade). The model to explain the neuronal firing rates was a linear, mixed-effects model while the one describing the binary, behavioral responses in those sessions was a binomial, generalized linear mixed-effects model. Both models were fit using Matlab’s *fitlme* function.

For both models, the fixed effects included the prior, saccade-driven noise, image noise, displacement, and interaction terms between image and saccade-driven noise, image noise and the prior, and saccade-driven noise and the prior. For the linear model predicting neuronal firing rates, we also included as random effects distinct intercepts and slopes for each of these predictors for each neuron. For the binomial model predicting behavioral responses, we included the same random effects but grouped by session rather than neuron identity.

### Code accessibility

- Neural, behavioral, and modeling datasets have been deposited at <u>Zenodo</u> and are publicly available at https://zenodo.org/records/14654929, affiliated with *bioRxiv* preprint https://doi.org/10.1101/2024.12.31.630950.
- All original code has been deposited at <u>GitHub</u> and is publicly available at https://github.com/dsub-neuro/FEF-paper-code.
- Any additional information required to reanalyze the data reported in this paper is available from the corresponding author upon request

## Results

### Behavioral prior use increased with saccade-driven noise but decreased with image noise

Two rhesus macaques (Monkey S and Monkey T, both males) performed a Saccadic Suppression of Displacement task (Bridgeman et al., 1975) in which they reported whether a visual stimulus in the periphery moved during an instructed saccade (Fig. 1c). Subjects first fixated a central square and then, on most trials, made a saccade to a target square. A Gaussian visual stimulus (“probe”), initially presented during fixation, was then displaced by varying amounts *during* the saccade. If the probe moved, subjects were required to report it with a second saccade to the probe. If it did not move, they were required to withhold their second saccade and maintain fixation on the target. Probe displacements were drawn from relatively broad and narrow Gaussian distributions in the movement (μ = 0º, σ = 2.5º) and non-movement (μ = 0º, σ = 0.2º) conditions, respectively. The instructed “intervening” saccade was always directed away from the recorded neuron’s receptive field (RF), and the probe always landed at the center of the post-saccadic RF.

We tested the Bayesian hypothesis that priors are used more with greater sensory uncertainty, either induced by the saccade itself or by noise in the external image (Fig. 1d). The prior probability of a probe jump (0.2 or 0.8) was indicated by the color of the fixation and target square. Two noise manipulations defined two experiments conducted on randomly interleaved trials: For the *image noise experiment*, the Gaussian probe had low, medium, or high noise as determined by its width, and all trials required an intervening saccade to the target. For the *saccade-driven noise experiment*, uncertainty was manipulated by either requiring an intervening saccade (high noise) or requiring steady fixation instead (low noise), all while a low image noise Gaussian probe moved. Critically, the two experiments had a condition in common: the low image noise condition with an intervening saccade occurred as a trial type for both.

We compared the monkeys’ behavior to the predictions of two models introduced in Subramanian et al., 2023, a Bayesian ideal observer model for all conditions (Fig. 2a-c) and a model combining a Bayesian component for saccade-driven noise and a perceptron-like category-learning network for image noise (Fig. 2d-f). Consistent with our previous findings, the monkey behavior tested here was consistent with the predictions of the combined model (Fig. 2g). We briefly describe the models below (detailed quantitative descriptions provided in *Methods* and Subramanian et al., 2023).

First, Bayesian ideal observer predictions were simulated as follows (Fig. 2a). On every trial, the ideal observer decides whether the probe “jumped” or not given the sensory evidence of a displacement, *x*. The decision is “yes” if the perceived displacement exceeds a threshold at which it was equally likely that the probe did or did not jump given its prior probability of jumping, and “no” otherwise. For each prior, *P*(*J*), the threshold is at a different location. For *P*(*J*) = 0.5, it was where the two likelihood distributions intersected, since each likelihood was weighted equally (Figure 2a, black vertical line). If there was no sensory uncertainty, the ideal observer would report “no jump” for all displacements less than the threshold and “jump” for all displacements above the threshold. Since the target was a Gaussian blob, we assume Gaussian uncertainty about the true displacement, *x* (Figure 2a, black distribution) and the decision was the integral over values of *x* greater than the decision threshold (Figure 2a, shaded region). This restricted the value of the decision to range from 0 to 1 and provided predictions that take the form of a psychometric function, i.e., the probability of reporting that the probe “jumped”, as a function of displacement.

For a high prior, e.g., *P*(*J*) = 0.8, the threshold would move to the left since *P*(*x*|*J*) was weighted higher than *P*(*x*|¬*J*) (the probability there was no jump), and vice versa for a lower prior, e.g. *P*(*J*) = 0.2. Critically, for the same displacement, the ideal observer was more likely to report that the probe jumped for a higher prior than for a lower prior, thus inducing a shift in psychometric curves across prior conditions (as seen in the blue and orange curves in Fig. 2b). We simulated increased saccade-driven noise as a widening of one’s internal non-jump likelihood distribution (i.e., *P*(*x*|¬*J*), consistent with the idea that it would take a larger displacement to be perceived across a saccade than during fixation. Image noise was incorporated in the model by widening the Gaussian uncertainty about the target, *σ*_*t*._ Taken together, the Bayesian model predicts that the shift in subjects’ psychometric curves across prior conditions – a measure of prior use – increases with both saccade-driven and image noise (Fig. 2c).

The combined model included a Bayesian component simulating saccade-driven noise (Fig. 2d, left) and a two-layer perceptron-like categorization network to predict behavior under increased image noise (Fig. 2d, right). The input layer of the categorization network consisted of units representing continuous displacements, with one set each per prior condition (orange and blue circles in Fig. 2d, right). The output layer had two units: “jump (J)” and “no jump (NJ).” Sensory noise was simulated as a Gaussian distribution of input unit activation, such that the total activation of input units was always 1. On each trial, the distribution was centered on the true displacement for the trial and the width of activation was determined by the sensory noise level. Each input unit was connected to both output units. The activation of each output unit was the weighted sum of inputs, and the final output of the perceptron network was normalized to be bounded between 0 and 1. To evaluate the possibility of crosstalk between the two noise types and model components, we fit the model to subjects’ responses using maximum likelihood estimation and allowed both noise parameters to vary freely across all trial types. The model fits were consistent with the separation of the two noise types across model components (Fig. S1).

We combined the outputs of the Bayesian and perceptron models at relative weights of 0.1 and 0.9, respectively, to generate the simulated data in Fig. 2e-f. Predicted outputs at additional Bayesian weights of 0.25, 0.5, 0.75, and 0.9 are shown in Figure S2. The combined model predicted that prior use increases with saccade-driven noise, as predicted by the Bayesian model, but *decreases* with increasing image noise in an “anti-Bayesian” manner (Fig. 2e; quantified in Fig. 2f). Consistent with these predictions and our previous report, subjects used priors more with the saccade-driven noise but less with increasing image noise in the current experiment even when the two trial types were interleaved (Fig. 2g). The difference in response rates across priors was greater for high compared to low saccade-driven noise (Monkey S: p=1.8×10^−6^ and Monkey T: p=9.32×10^−4^; paired t-tests). However, it decreased with greater image noise (Monkey S: F(2) = 50.5; p=1.1×10^−14^ on a repeated-measures ANOVA with p=5.04×10^−6^ for low vs. medium noise, p=1.60×10^−10^ for low vs. high noise, and p=4.74×10^−7^ for medium vs. high noise; Monkey T: F(2) =8.67, p=0.001 on a repeated-measures ANOVA with p=5.38×10^−4^ for low vs. medium noise, p=0.01 for low vs. high noise, and no significant difference between medium and high noise (p=0.27)). A comprehensive characterization of psychometric curves from the same animals is shown in Subramanian et al. (2023) and the raw responses across prior and noise conditions for the sessions used here shown in Fig. S3. In summary, behavior was consistent with Bayesian predictions when faced with saccade-driven noise but was anti-Bayesian for image noise, instead reflecting a flexible, prior-dependent adjustment of a category boundary.

### FEF firing rates reflected the prior, but only when the sensory evidence was available

We then asked if FEF neurons reflected the priors and if its activity predicted either the Bayesian or categorization behaviors. We first assessed whether the activity of FEF neurons reflected the priors. Analyses were based on 79 FEF neurons for which there were at least 10 trials completed in each task condition (53 from Monkey S, 26 from Monkey T). We focused on the common condition (low image noise with an intervening saccade) in which behavioral prior use was the highest (Fig. 2g). We pooled across displacements to maximize statistical power and ensure approximately equal numbers of trials across the prior conditions.

As shown in Figure 3a, population firing rates in the low (0.2, teal) and high (0.8, orange) prior conditions were different. Yet this difference did not emerge when the prior for the trial was first revealed to the animal, but only when the sensory evidence about the displacement entered the RF after the intervening saccade. This type of post-saccadic sensory stimulation of the RF is called *reafference*. The prior-modulated firing rate difference persisted from the start of the reafferent stimulation, past the median saccadic response time (611 ms), and then converged back together around 800 ms after the reafferent event. Note, since FEF activity is sensitive to the magnitude of the displacement, and displacements were unequally represented across the two prior conditions, we also confirmed that the results were qualitatively similar even when displacement was 0 (Fig. S4).

**Figure 3.**
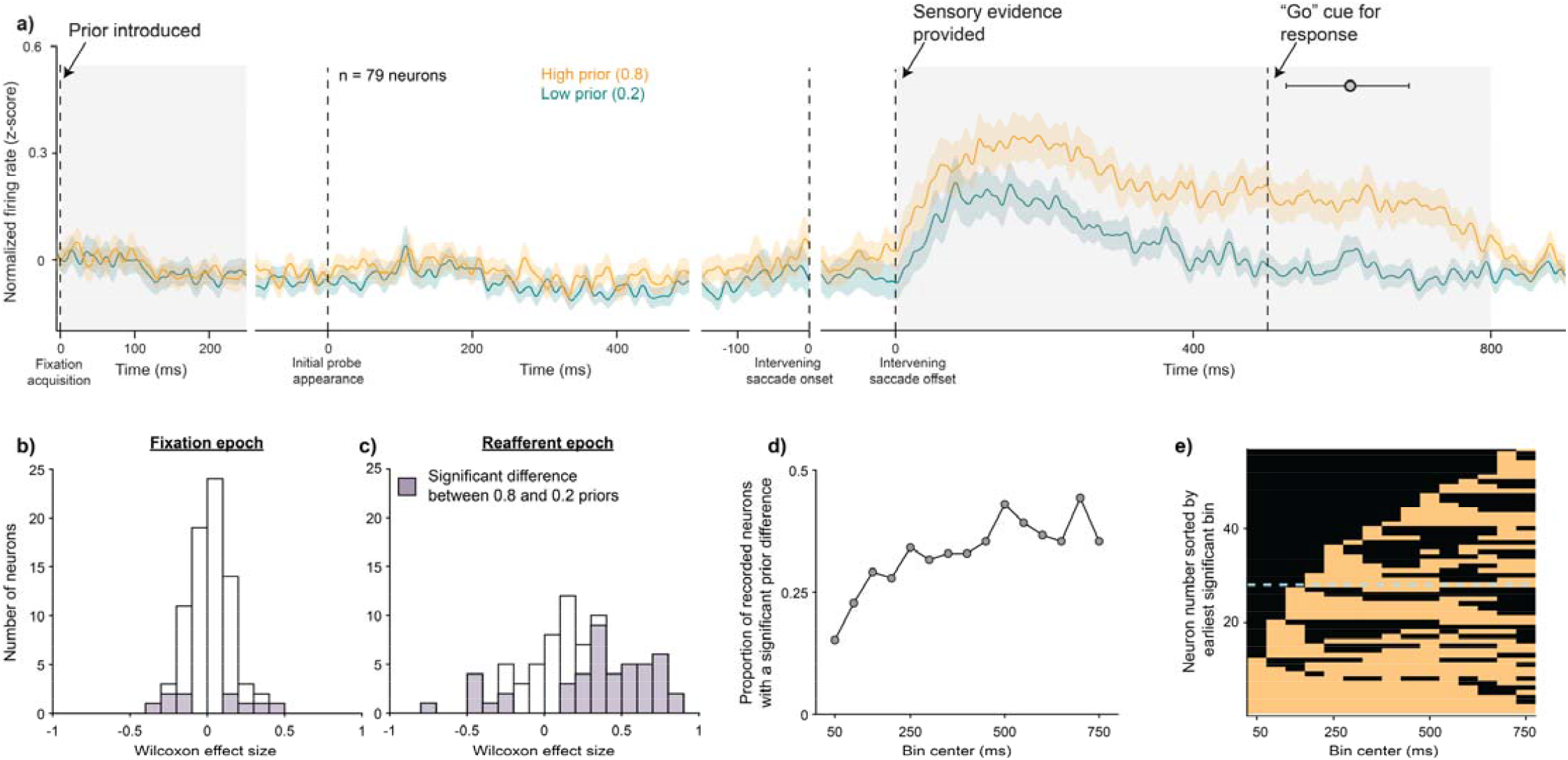
FEF activity reflected the prior but only when the sensory evidence was presented. Data are from the common condition (low image noise with an intervening saccade). a) Normalized peri-stimulus time histograms averaged across all 79 neurons aligned to various task events. A clear difference between the high (orange) and low (teal) prior responses emerged not when the prior was initially introduced at fixation acquisition but only when the sensory evidence for the task (i.e., the post-saccadic location of the probe) was presented. Gray circle at 611 ms indicates the median response time on trials where a jump was reported with a saccade (error bars range from the 25^th^ to 75^th^ quartiles). b-c) Histograms of Wilcoxon effect sizes between the high and low prior conditions in the fixation (0-250 ms from fixation acquisition) and reafferent (0 – 800 ms from the intervening saccade offset) epochs. Cells with significant differences between the prior conditions (p < 0.025 to correct for two comparisons) are shown in purple. d) Same neural dataset as in panel c but shown as the proportion of total neurons with significant prior effects during each overlapping 100 ms bin of the reafferent epoch (p < 0.00167 to correct for 15 comparisons). e) Same neurons again but ordered by the earliest significant bin. Significant prior effects are colored orange. Almost half of the significant cells (dashed blue line) showed a prior effect within 150 ms of the reafferent event.

To quantify this prior-modulated activity, we focused on two task epochs (highlighted in gray in Fig. 3a) during which the spatial locations of the fixation square, target square, and Gaussian probe relative to the RF were the same across all task conditions (Fig. 1d). While only 12.6% of the neurons (10/79) showed a statistically significant prior selectivity in the fixation epoch (0-250 ms from fixation acquisition, Fig. 3b), over half the neurons (58.2%, 46/79) were selective in the post-jump epoch (0-800 ms from the reafferent event, Fig. 3c; significant effect sizes, shaded, were evaluated using Wilcoxon signed-rank tests with α = 0.025 to correct for two comparisons). Distributions for each animal are shown in Figure S5. We analyzed the time course of the effect in the post-jump epoch using successive 100 ms bins (with 50 ms overlaps). This showed that 72.2% of the neurons (57/79) were significantly selective during at least one 100 ms bin (Fig. 3d; Wilcoxon signed rank tests with α = 0.00167 (0.025/15) to further correct for 15 comparisons), and many neurons remained persistently selective rather than exhibiting a clear sequencing of activity across neurons (Fig. 3e). In summary, FEF neurons reflected the prior, but only after the sensory evidence was available. We thus focused our subsequent analyses on this reafferent period of the task.

### FEF neurons predicted anti-Bayesian but not Bayesian prior use

Next, we asked how the prior modulation identified in the common condition (second column of all rows in Fig. 4) was modulated by saccade-driven or image noise. Possible outcomes were that the prior signal might be modulated 1) only by saccade-driven noise but not image noise (i.e., Bayesian, Fig. 4a); 2) only by image noise but not saccade-driven noise (anti-Bayesian, Fig. 4b); 3) by neither the image nor the saccade-driven noise (Fig. 4c), suggesting that FEF neurons report the prior but do not reflect either computational strategy for visual stability; or 4) by both the image and saccade-driven noise (Fig. 4d), suggesting that FEF neurons do not differentiate between the computations but simply reflect the final behavioral output.

**Figure 4.**
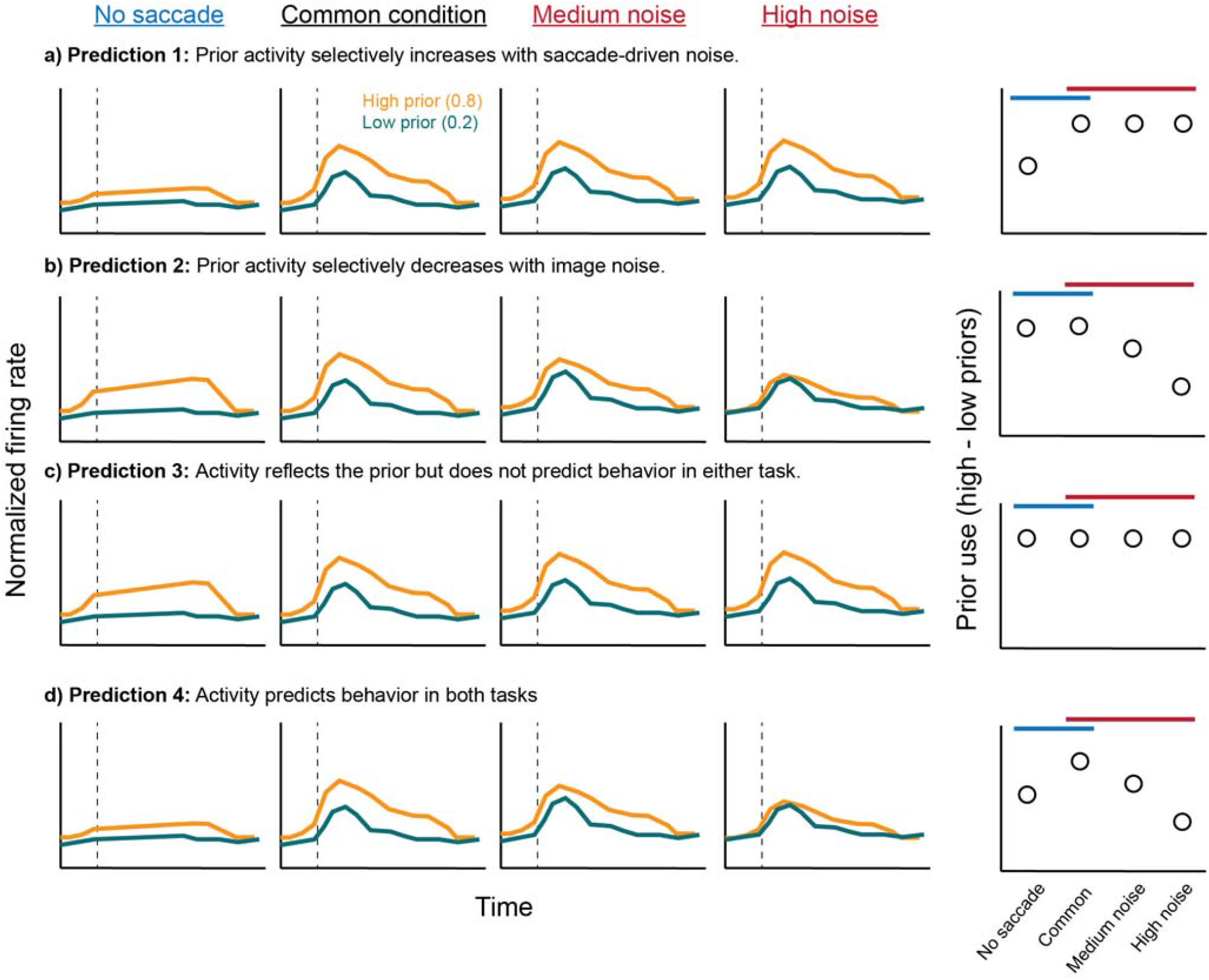
Predicted outcomes for prior use across the four noise conditions. Each row shows a different prediction. Columns 1-4 show cartoons of peristimulus time histograms in the high (orange) and low (teal) prior conditions, aligned to the reafferent event (dashed line), for the four noise conditions across the columns (see label above each column). The second column depicts the common condition (with saccade and low image noise, same across all predictions). The fifth column summarizes the prior difference across the four noise conditions.

The results most closely matched the second possible outcome (Fig. 4b): the prior signal was not significantly modulated by saccade-driven noise but, instead, only by image noise (Fig. 5). The difference in firing rates between priors did not exhibit a qualitative change between the no-saccade (Fig. 5a) and the saccade condition, which was also the common condition (Fig. 5b). However, the difference in firing rates between priors decreased from the common condition, which used low image noise, to the medium (Fig. 5c) and high image noise (Fig. 5d) conditions. As in the case of the prior effects, we also pooled across displacements here to maximize statistical power and ensure an equal distribution of trials across priors. We confirmed, however, that the same effect was found with the displacement restricted to 0 (Fig. S6). Statistically, the Wilcoxon effect sizes of the difference between priors across all 79 neurons were not significantly different between the no-saccade condition and the common condition (p = 0.81, paired t-test), but they were significantly lower than the common condition in the medium (p = 0.013) and high (p = 0.0047) image noise conditions (Fig. 5e).

**Figure 5.**
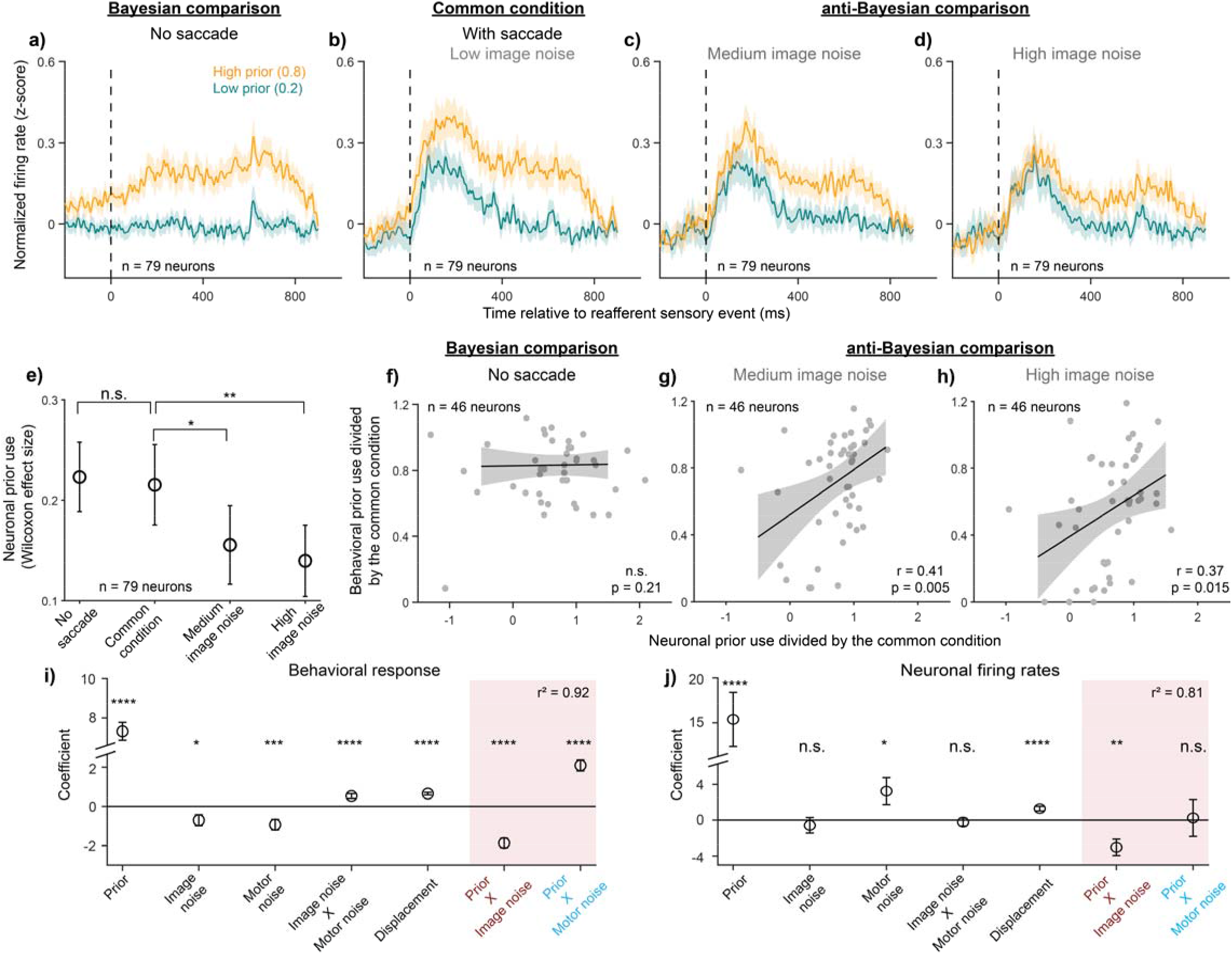
FEF neurons predicted the anti-Bayesian but not Bayesian behavior. a-d) Peristimulus time histograms in the high (orange) and low (teal) prior conditions, aligned to the reafferent event (dashed line), for the a) no-saccade, b) common condition with saccade and low image noise, c) with saccade and medium image noise, and d) with saccade and high image noise conditions. Comparison between the no-saccade condition and common condition allows for the evaluation of whether FEF activity reflects Bayesian behavior, while comparison between the common, medium image noise, and high image noise conditions allows for evaluation of whether activity reflects the anti-Bayesian behavior. e) Wilcoxon effect sizes between the high and low prior conditions across the four noise conditions. f-h) Session-by-session correlation between high vs. low prior Wilcoxon effect sizes (i.e., neuronal prior use) and behavioral prior use for the f) no-saccade, g) medium image noise, and h) high image noise conditions, when all values were normalized to the common condition. i) Coefficients from a mixed-effects, binomial generalized linear regression predicting the binary behavioral response (“jumped” or “did not jump”). j) Coefficients from a mixed-effects, linear regression predicting firing rates. *, p < 0.05, **, p < 0.01, ***, p < 0.001, ****, p < 0.0001. Data in f-j include only the neurons, and their corresponding behavioral sessions, that had significant prior selectivity in the common condition (n = 46 neurons, evaluated using a Wilcoxon signed-rank test).

For neurons with significant prior selectivity in the common condition (i.e., p < 0.025, corrected for comparisons in the fixation and post-jump epochs), we further found that neuronal prior use predicted behavioral prior use session-by-session in the image noise condition but not the saccade-driven noise condition. To directly compare the change in neuronal and behavioral prior use with the two uncertainty types within each session, we normalized both to the common condition for that session. The correlation between the normalized neuronal and behavioral prior use was not significant for the no-saccade condition (Fig. 5f, p = 0.21) but was significant with a positive correlation for the medium (Fig. 5g) and high (Fig. 5h) image noise conditions (r = 0.41, p = 0.0050 for medium noise and r = 0.37, p = 0.0151 for high noise; Pearson correlations).

We additionally considered that for all conditions with an intervening saccade, the probe was initially placed near the RF center *after* the saccade, but for the no-saccade condition, it was placed near the pre-saccadic RF center since there was no movement of the RF (Fig. 1d, leftmost column). Since displacements were typically smaller than 5°, it was likely that the initial location of the probe was already within the RF in this condition (Mayo et al., 2015). Therefore, it was possible that the prior selectivity and its interaction with behavior aligned better to the initial probe presentation than to the probe jump event (analogous to a reafferent response) for the no-saccade condition. However, while there was clear transient visual activity aligned to probe presentation near the RF center as expected, the neurons showed no prior selectivity during the initial probe presentation epoch, and we did not find a significant correlation between neuronal prior use and behavior (Fig. S7).

Finally, to examine the possibility that other experimental variables (magnitude of displacement, image noise levels, the presence of an intervening saccade, and the generation of a saccade into the neurons’ RFs) may have had confounding effects on the correlation tests of Figure 5f-h, we performed regression analyses on the data. One major consideration was the generation of saccades into the RF since FEF activity is known to be modulated by saccades. However, since the saccade into the RF was also the animal’s response on the trial, all other experimental variables such as displacement magnitude, priors, noise levels, etc. were correlated with it. Thus, including both sets of variables as predictors for the same regression model was not possible. To get around this issue of multicollinearity, we ran two multiple, mixed-effects regression models and compared their coefficients. The first was a logistic model which used the *binary behavioral responses* of the animals as the regression model outcome (Fig. 5i). The other was a linear model which used the *firing rates of the neurons* as the outcome (Fig. 5j; detailed descriptions in Methods). All experimental variables significantly predicted the behavioral response, as expected (Fig. 5i), including the interactions between priors and both image (negative coefficient, the anti-Bayesian effect) and saccade-driven noise (positive coefficient, the Bayesian effect; red shading). When predicting the neural firing rates, however, only the interaction between priors and image noise had a significant effect (negative), while the interaction between priors and saccade-driven noise did not (Fig. 5j, red shading). Three other variables had a significant effect on firing rates, too, all of which were expected. Priors had a strong positive effect since only the cells with a significant prior effect were included. Additionally, saccade-driven noise (i.e., the presence of an intervening saccade) and displacement had significant positive coefficients indicating that FEF activity is sensitive both to the presence of an image at the center of the post-saccadic receptive field and the amount by which it moved there. These results were consistent even when all recorded neurons were modeled (Fig. S8). Taken together, although the animals’ behavioral prior use was modulated *both* by saccade-driven and image noise, the prior signal in FEF was modulated *only* by image noise. That is, FEF activity selectively correlated with and predicted only the “anti-Bayesian” prior use, consistent with the use of a categorization model.

## Discussion

While we previously showed that judgments of visual stability across saccades exhibit both Bayesian and anti-Bayesian effects of noise from saccadic and visual sources, respectively, here we found that FEF neural activity is significantly modulated only by the latter. This suggests that the two behavioral strategies are implemented by distinct circuits in the brain and that FEF is part of a pathway that selectively implements the anti-Bayesian behavior. Indeed, the timing of the observed prior signal in FEF, i.e., its emergence only at the reafferent event rather than when the prior is first introduced, is consistent with the dissociation between Bayesian inference and our proposed categorization model in FEF. For Bayesian inference, the prior and sensory evidence are eventually combined into a posterior for a decision. However, they must be encoded independently at some stage of processing such that uncertainty in one of the factors does not affect the other. We do not see such independent encoding of priors and sensory evidence in FEF. Additionally, we proposed in Subramanian et al. (2023) and recapitulated in Fig. 2. that decreasing prior use could be explained by a perceptron-like, two-layer neural network that associates the continuous sensory input (displacement amounts) with categorical “jumped’ or “did not jump” response outputs. A critical assumption was that the association (continuous to categorical) was dependent on the context provided by the prior information, leading to a difference across prior conditions in the response rates at each displacement. When external noise is added to the sensory evidence, it blurs the input-output associations across prior contexts such that they appear to collapse together, both in behavior and, as shown here, neuronal activity. In this sense, our findings that prior and image noise effects emerge at the reafferent event and span the time frame until the response is made are both consistent with a representation of the sensory input about displacements being associated with the response. The FEF’s involvement in categorization in our task is also consistent with the proposed function of the frontal orienting fields (FOF), an analogous region in the rodent brain, during perceptual categorization (Erlich et al., 2015).

These results raise the complementary question of where in the brain the Bayesian compensation for saccadic motor noise may be implemented. One possibility is the superior colliculus (SC). Neurons in the intermediate layers of the SC, an origin of saccadic corollary discharge signals (Sommer and Wurtz, 2008), have also been shown to signal probabilistic priors for binary sensory decisions (Crapse et al., 2018; Jun et al., 2021) and neurons in superficial SC are modulated by saccadic suppression (Walker et al., 1995). The SC thus signals both necessary components of the Bayesian computation for our task. However, the neural signatures of saccadic suppression have also been observed in a variety of regions in the visual system including the Medial Temporal region (MT), the Medial Superior Temporal region (MST), the Ventral Intraparietal region (VIP) and the Lateral Intraparietal region (LIP) (Bremmer et al., 2009; Thiele et al., 2002), area V4 (Zanos et al., 2016), and even in the retina (Idrees et al., 2020). The possible contributions of these regions, along with the SC and FEF, to a distributed circuit for visual stability remain to be clarified. Our results, showing the use of two distinct computational strategies as well as their dissociation in the brain, provide a first step towards an integrated understanding of the behavioral, computational, and neural circuit basis for perception across saccades.

We consider a few additional points in interpreting the results. First, in addition to visual responses, many neurons in FEF exhibit activity synchronized to saccades (Bruce and Goldberg, 1985; Dias and Segraves, 1999; Robinson and Fuchs, 1969, Sommer and Tehovnik, 1997). Since response saccades were made into neurons’ RFs in our task, it is likely that at least part of the prior modulation we observed is due to the higher proportion of response saccades in the high prior condition. Indeed, individual neurons’ choice probabilities (Britten et al., 1996) related to whether the animal reported “jumped” or not were highly correlated with their prior modulation across all noise conditions (Fig. S9, top row). However, saccade-related modulation is not sufficient to explain our results. Importantly, response saccades were made into a neurons’ RF for all task conditions, whether uncertainty was introduced by image noise or saccade-driven noise (Fig. 2d). Indeed, we restricted our analyses to epochs where the spatial configuration of the task was not different between the image and saccade-driven noise experiments. If the observed activity were driven solely by the response saccade, then it would simply be correlated with the output behavior in all conditions (as predicted in Fig. 4d) rather than displaying a dissociation between non-Bayesian and Bayesian behavior across noise types. Consistent with this, neuronal choice probabilities related to “jump” vs. “non-jump” decisions did not show strong correlations with animals’ behavioral prior use on a neuron-by-neuron basis (Fig. S9, bottom row). Additionally, while the median response time was 611 ms, the prior difference and its modulation by image but not saccade-driven noise emerged as early as 0-100 ms after the reafferent sensory visual event with about half the modulated neurons displaying significant prior differences within 200ms (Fig. 3e), further suggesting that the dissociation between computations reflected in FEF starts with the sensory event rather simply reflecting the motor response. Finally, our regression analyses quantitatively show that while the saccades (i.e., subjects’ responses) were influenced by both image and saccade-driven noise, the neurons’ firing rates were influenced only by image noise. A task design that explicitly dissociates the sensory information from the reporting action may be used to address this question more definitively in the future. Additionally, FEF activity is also influenced by other more cognitive factors such as decision confidence (Fan et al., 2024) or attentional gain (Squire et al., 2013), and our results may reflect a modulation by these factors as well. For example, a higher expectation of probe movement in the high prior condition may lead to a higher confidence response, and addition of noise may cause a collapse of that confidence. Similarly, the high prior may lead to the deployment of greater attention to the probe and this attentional effect may may also be modulated by noise. While neural activity in the image noise conditions may be consistent with these factors, activity in the saccade noise conditions are not. For saccade noise trials, prior modulation of activity shows no significant change between the no-saccade and with-saccade, low noise conditions. If decision confidence or attention were strong modulators of our neural activity, one would expect them to have the same effects in the two types of interleaved noise trials, but they don’t.

Second is the question of the scope of the dissociation between Bayesian and discriminative processing in FEF. Is there any context in which the activity of FEF neurons may be consistent with the implementation of Bayesian inference? We and others have found that humans are Bayesian when asked to report the *continuous* displacement of a probe rather than providing a categorical report of whether it jumped or not (Subramanian et al., 2023; Niemeier et al., 2003). Visual receptive fields in the FEF, both before and after remapping, maintain their spatially continuous retinotopic organization, and FEF neurons are continuously tuned to the magnitude of image displacements across saccades (Crapse and Sommer, 2012). It is possible, therefore, that the FEF supports the implementation of a continuous Bayesian model for visual stability across saccades, as it does in other sensory (Langlois et al., 2025) and motor (Darlington et al., 2018) contexts. A direct test of this possibility is feasible but has not been done. If indeed FEF activity conformed with the Bayesian prediction in this case, this would suggest that FEF supports Bayes optimal estimates when the stimuli match how visual location information is intrinsically organized within the area, i.e., in continuous rather than categorical coordinates.

Third, we consider the implications of our results for interpreting the role of Bayesian inference in sensory processing. The success of the Bayesian framework in explaining behavior has spurred a search for how Bayesian computations may be implemented by the brain. Leading theoretical proposals include the ideas of “Probabilistic Population Codes,” in which sensory variability automatically represents the likelihood and prior distributions required for Bayesian integration (Ma et al., 2006; Beck et al., 2008); of Bayesian inference via efficient coding, in which priors are inherently represented by the matched allocation of neural infrastructure to the natural statistics of the environment (Wei and Stocker, 2012; Ganguli and Simoncelli, 2014; Park and Pillow, 2024); and of predictive coding models, in which feedback projections convey prior information that is integrated with feedforward sensory likelihoods (Aitchison and Lengyel, 2017). These proposals have in common the intuition that Bayesian inference in simple sensory contexts results from the natural, evolutionarily shaped functioning of sensory circuitry rather than reflecting an explicit, reward-maximizing strategy that needs to be effortfully generated. If this were true, Bayesian inference stands to serve as a general computational principle of sensory function (Knill and Pouget, 2004; Friston, 2012). Our results showing that the reafferent visual responses in FEF are not consistent with Bayesian ideal observer predictions, even in a context where the behavior *is*, caution that Bayesian inference is not the default mode of sensory circuits, but that the precise relationship between Bayesian behavior and sensory processing should be examined in a context-dependent manner.

Finally, our results have implications beyond the context of visual stability. All sensory systems must account for self-movement to maintain coherent percepts in naturalistic contexts. It is possible that our observed dissociation between computational strategies in the face of self-generated vs. external uncertainty extends to other perceptual domains as well. Additionally, Bayesian models are an example of optimal, generative models, and the category-learning we proposed is an example of a simpler heuristic. Although there is growing interest in the relative contributions of optimal vs. heuristic models in sensory processing (Carandini, 2024; Gardner, 2019, Laquitaine and Gardner, 2018; Rahnev and Denison, 2018; Sohn and Jazayeri, 2021), Bayesian inference remains predominant in explaining performance in simpler stimulus estimation tasks. However, the two types of models are often used in conjunction in more cognitive contexts such as decision-making (Tversky and Kahneman, 1974; Gigerenzer and Gaissmaier, 2011). Our results addressing the neural basis for how Bayesian and discriminative models may be implemented for visual perception could help to serve as a bridge between our understanding of lower-level sensory processes and decision-making in higher order cognitive contexts.

## Acknowledgments

We thank Jessi Cruger for helping with the care of the non-human primates and Drs. Stephen Lisberger, Jeff Beck, and Greg Field for their insights, feedback, and discussions on the work. Supported by a 2020 Incubator Award from the Duke Institute for Brain Sciences, Duke University, to J.M.P. and M.A.S. Author D.S. is presently at Laboratory of Sensorimotor Research, National Eye Institute, National Institutes of Health, Bethesda, MD, USA, 20894.

## Author contributions

Conceptualization: D.S, J.M.P., and M.A.S., Methodology: D.S., J.M.P., and M.A.S.; Software: D.S.; Validation: D.S.; Formal Analysis: D.S. and J.M.P.; Investigation: D.S.; Resources: M.A.S.; Data Curation: D.S.; Writing – Original Draft: D.S.; Writing – Review and Editing: D.S., J.M.P., and M.A.S.; Visualization: D.S.; Supervision: M.A.S.; Project Administration: M.A.S.; Funding Acquisition: J.M.P., M.A.S.

## Supplemental information

**Figure S1:**
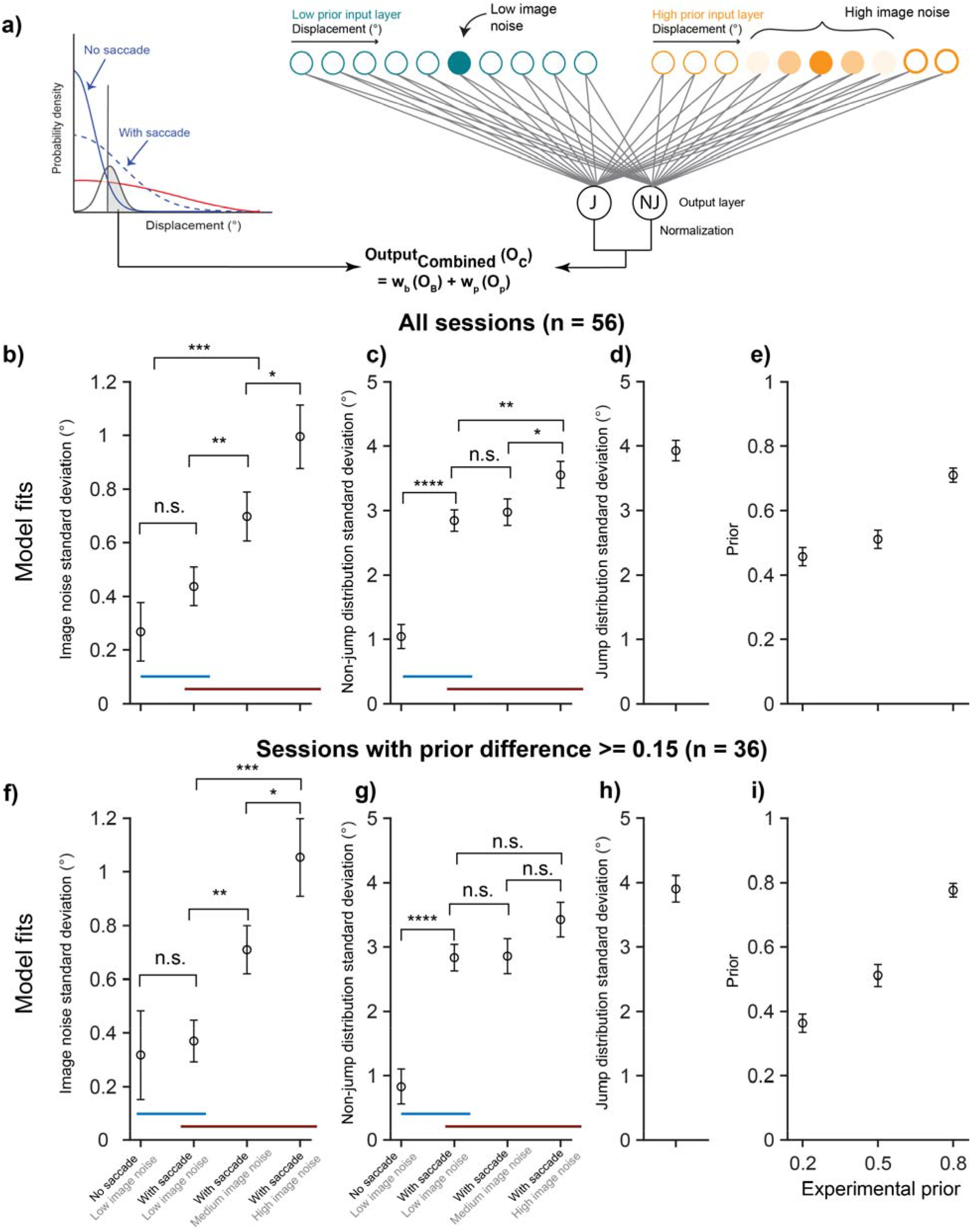
Combined model parameters fit to behavior. a) Schematic of the combined model (duplicated from Fig. 2d). Image noise is parametrized as the width of the distribution centered on the probe displacement and saccade noise is parametrized as the width of the “non-jump” displacement distribution centered on 0. Additionally, we also allowed the “jump” distribution and 3 levels of priors to vary as free parameters. Note that the two conditions with low image noise (with and no saccade) also included a 0.5 prior condition as in Subramanian et al., 2023. The data are not shown in this manuscript to maintain the same prior comparison between the 0.8 and 0.2 conditions across all noise levels, but those trials were included for model fitting. We fixed the weight of the Bayesian component at 0.1 as in the simulations. We fit the model separately to data from each session using maximum likelihood estimation. To simulate the learned weights between the displacement inputs and the “jump” and “non-jump” outputs for each prior condition, we used the value of the psychometric curve fit to all data in that prior condition for each session. b-e) Means and standard errors of fits across all sessions. The image noise parameter varies primarily across the image noise conditions (F(2) = 10.8; p = 5.2169e-05 on a repeated-measures ANOVA) but does not differ significantly between the no-saccade and common conditions (b). The model fits for the saccade noise parameter (i.e., the width of the non-jump distribution) vary significantly between the no-saccade and common conditions (p = 2.9514e-10 on a paired t-test; Cohen’s d = −1.3459) but much less across the image noise conditions (high image noise vs. low Cohen’s d = 0.51; high image noise vs medium Cohen’s d = 0.38; panel (c)). The model fits for the jump distribution (d) and priors (e) were consistent with expectations. f – i) Model fits for the priors themselves may trade off with the noise parameters, which primarily serve to modulate prior use in our model. To test this possibility, we evaluated model fits only for the sessions in which the fits for the low and high prior differed by at least 0.15 (n = 36 sessions). Indeed, for these sessions, there was a complete statistical dissociation between the saccade and image noise experiments. There was no significant difference between the image noise fits in the saccade experiment and significant variation across the image noise conditions (F(2) = 12.19; p = 2.8583e-05; (f)). Saccade noise fits varied significantly across the saccade noise conditions (p = 6.8431e-07) but not across the image noise conditions (F(2) = 2.28; p = 0.11; (g)). The model fits for the jump distribution (h) and priors (i) were once again consistent with expectations.

**Figure S2.**
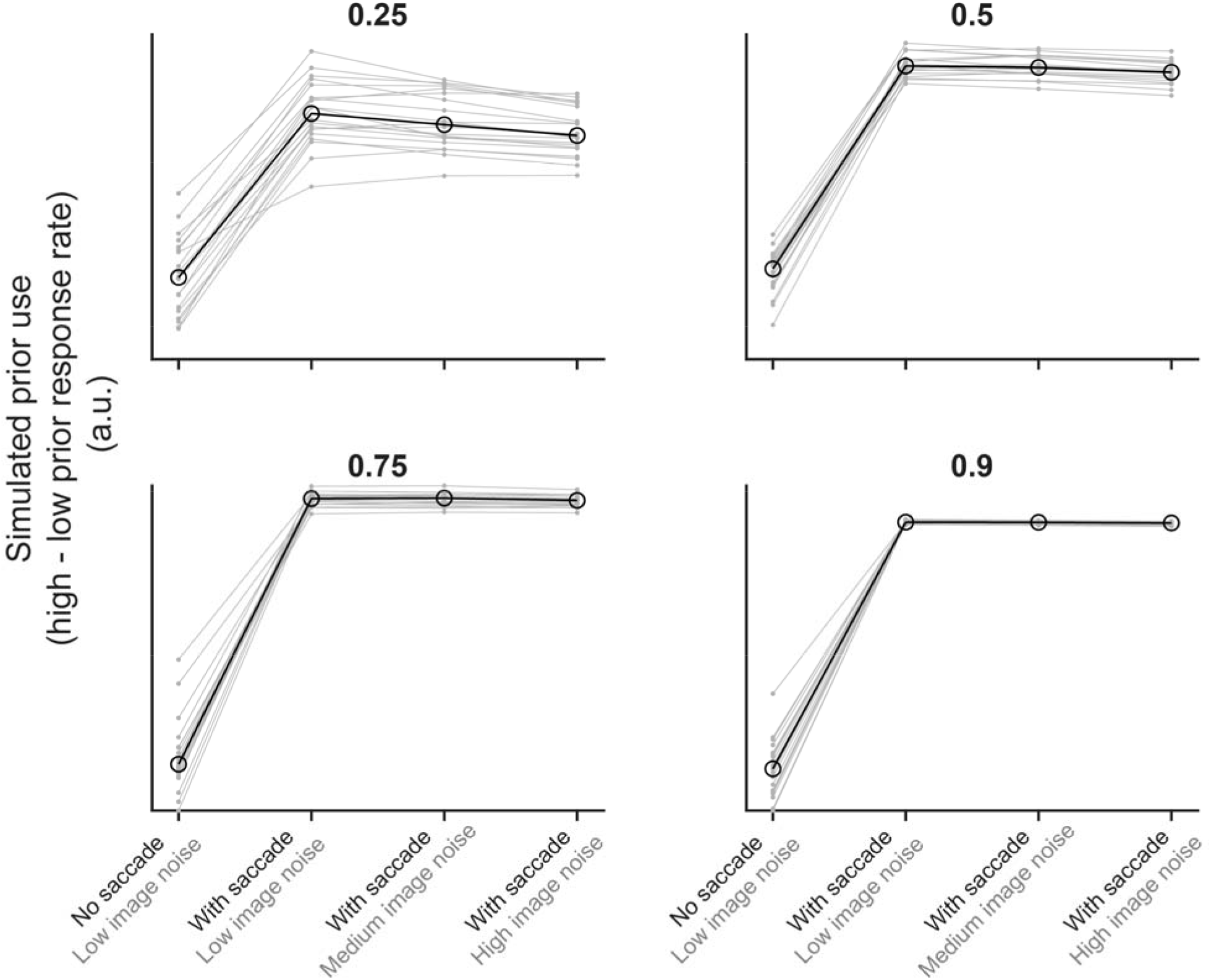
Predicted output of a Bayesian + classifier combined model (same as Fig. 2f) but at four additional weights of the Bayesian component (top left, 0.25; top right, 0. 5; bottom left, 0.75; bottom right, 0.9). Increasing the weight of the Bayesian component has two main effects. First, it modulates the extent to which prior use is influenced by saccade vs. image noise (i.e., slope of the line joining the leftmost two points vs the line joining the three points on the right). For the saccade noise component, this is expected since it is varied in the Bayesian component of the model. For the effect on the image noise conditions where saccade noise is high, the intuition is that the Bayesian component pushes the psychometric curves across prior conditions apart from each other but additional image noise in the categorization component pushes them together. If the weight of the Bayesian component is high enough, it pushes the curve separation to a ceiling such that the dynamic range of the effect of image noise is masked. Note that it does not switch to being a positive slope since image noise is not being varied within the Bayesian component. It reduces the variability in the simulated data. This is because the Bayesian component of the model is deterministic (i.e., for the same input parameter values, it will always produce the same output probability of reporting “jumped”), but the categorization component is not. Since “jumps” and “non-jumps” are drawn from probabilistic distributions and the feedback provided to the perceptron is contingent upon the distribution they were drawn from, it is possible for the same displacement to be classified as either a “jump” or “non-jump” on distinct iterations of the model.

**Figure S3.**
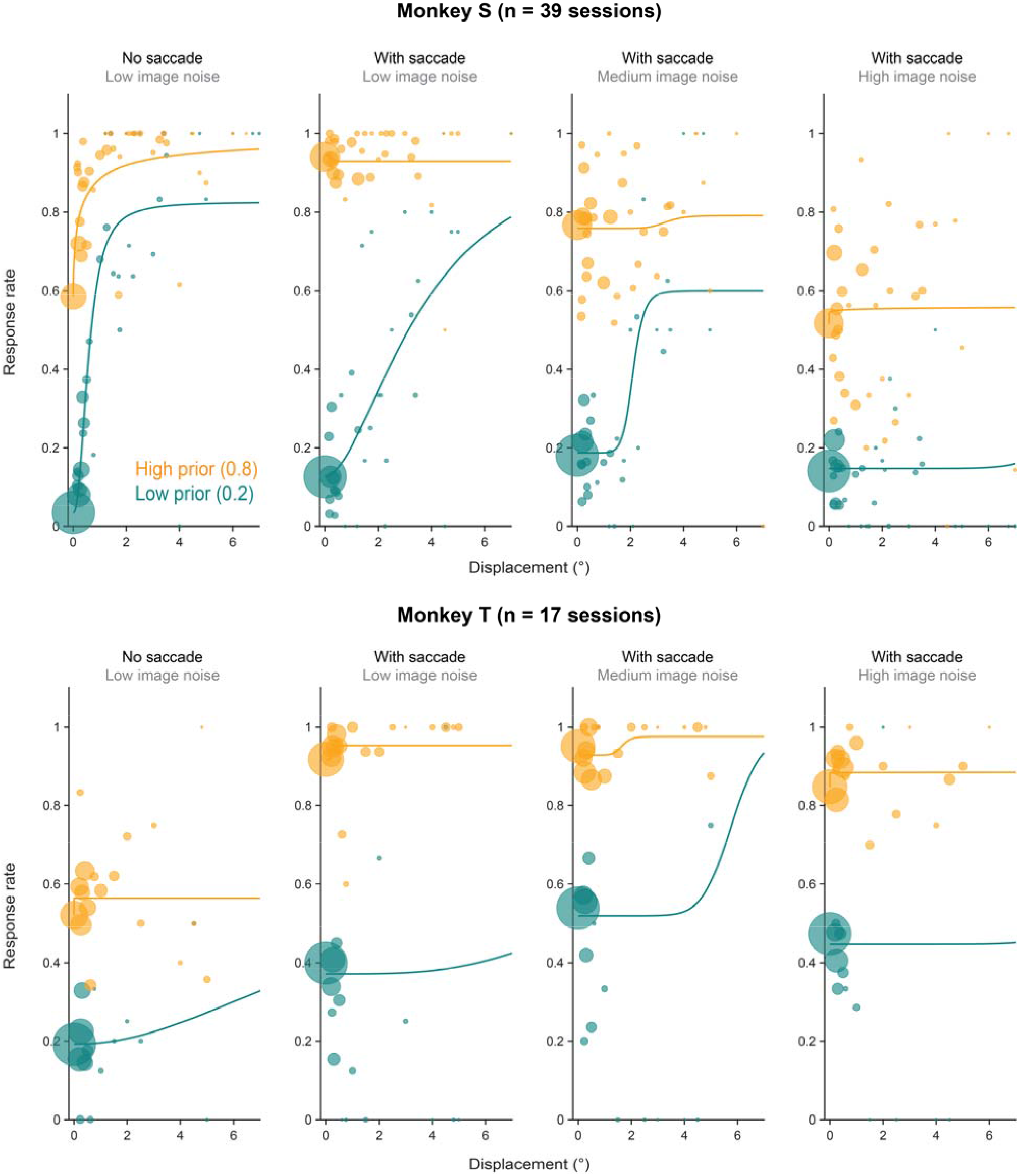
Response rates as a function of displacement across the prior and noise conditions for Monkey S (top) and Monkey T (bottom). Relative sizes of the bubbles indicate the relative numbers of trials at each displacement. Data are pooled across sessions for both animals. Lines show psychometric fits to the pooled data for ease of visualization, but we caution against interpreting their parameters since the spatial configuration and eccentricity of the probe were not held constant across sessions. A comprehensive characterization of psychometric curves for the same animals in versions of the image- and saccade-noise tasks with matched spatial parameters are shown in Subramanian et al. (2023).

**Figure S4.**
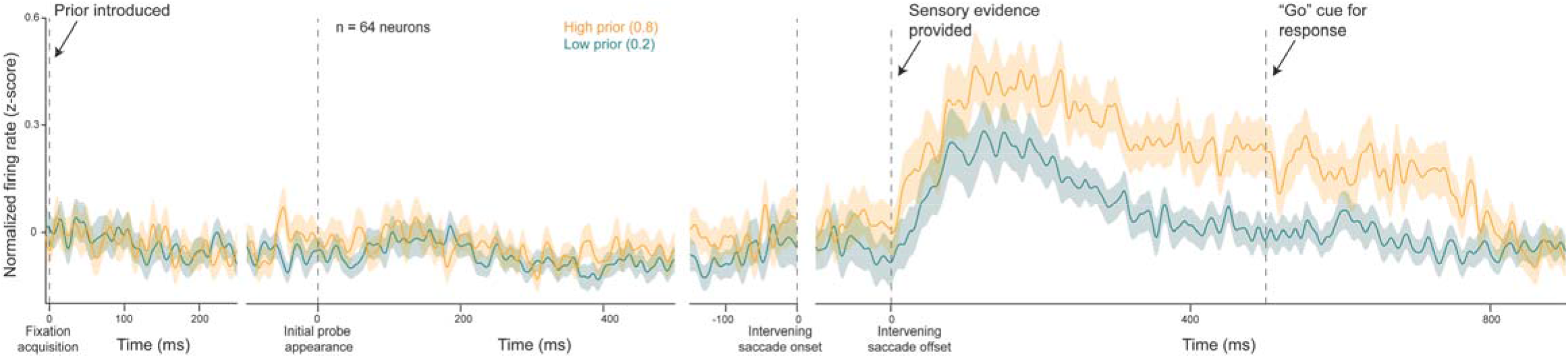
Same as Fig. 3a but restricted to trials where the displacement was 0.

**Figure S5.**
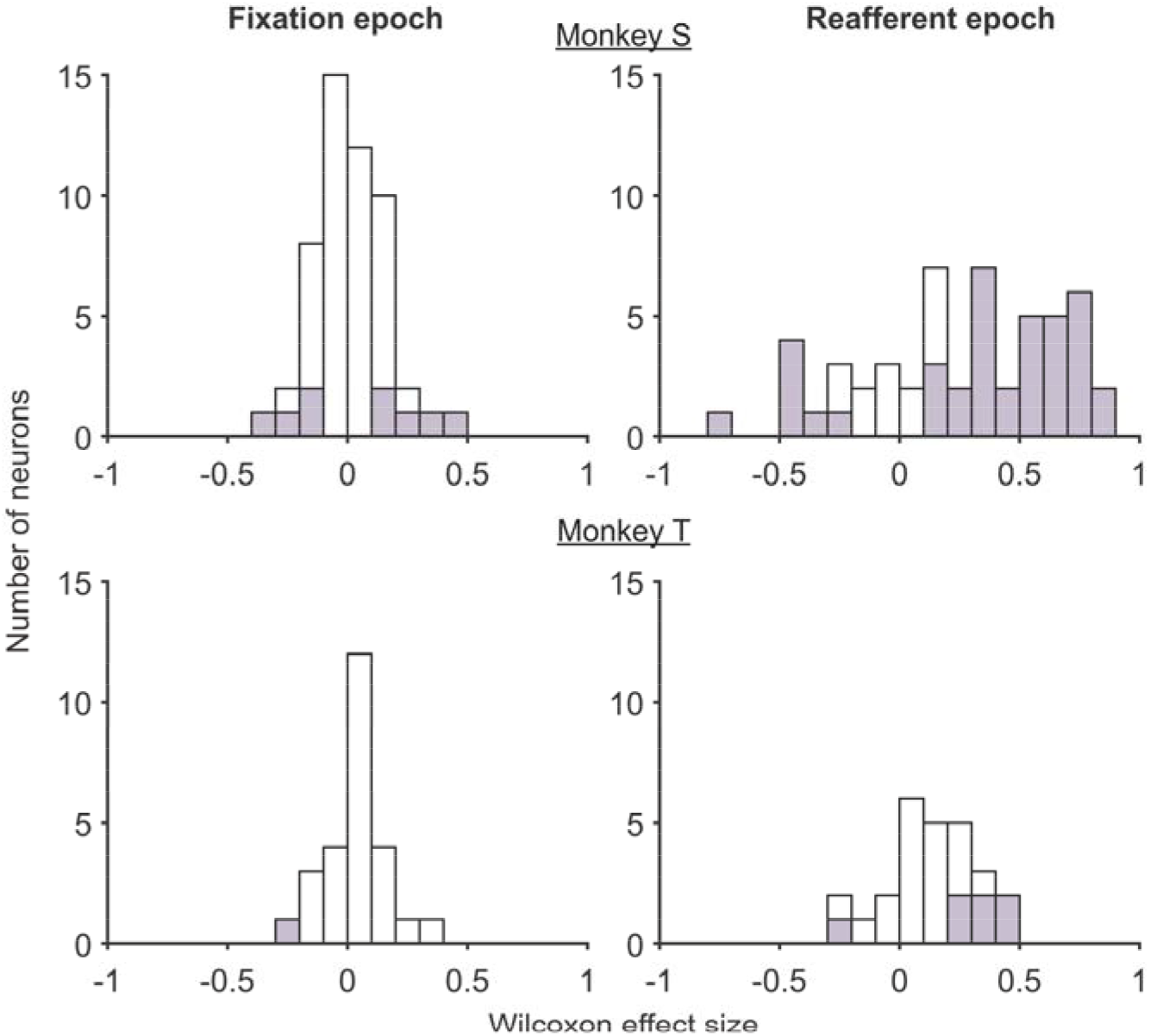
Same as Fig. 3b-c but for each animal individually. Top row: Monkey S, bottom row: Monkey T.

**Figure S6.**
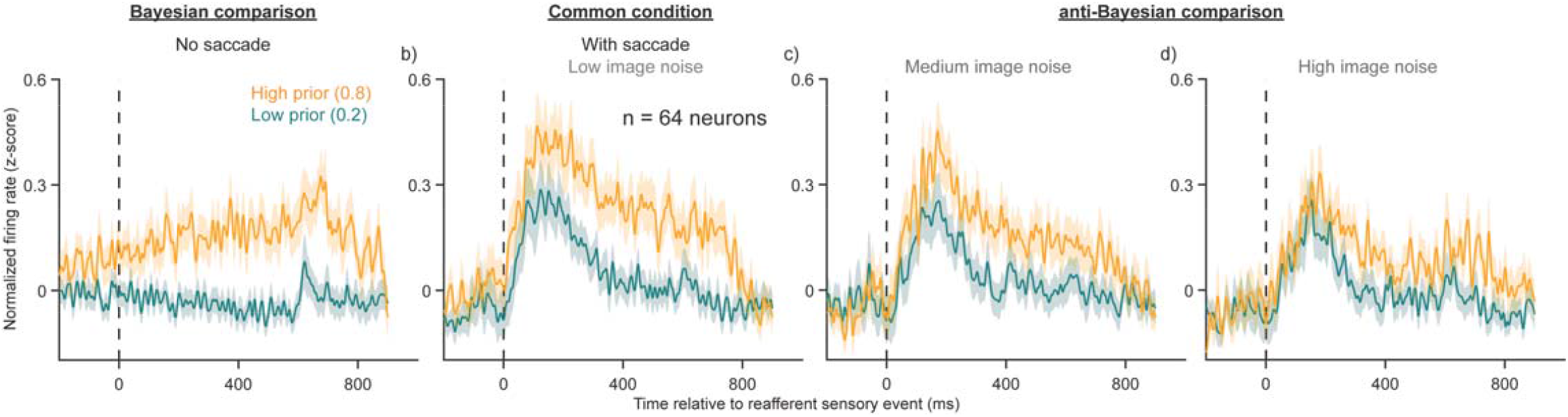
Same as Fig. 5a-d but restricted to trials where the displacement was 0.

**Figure S7.**
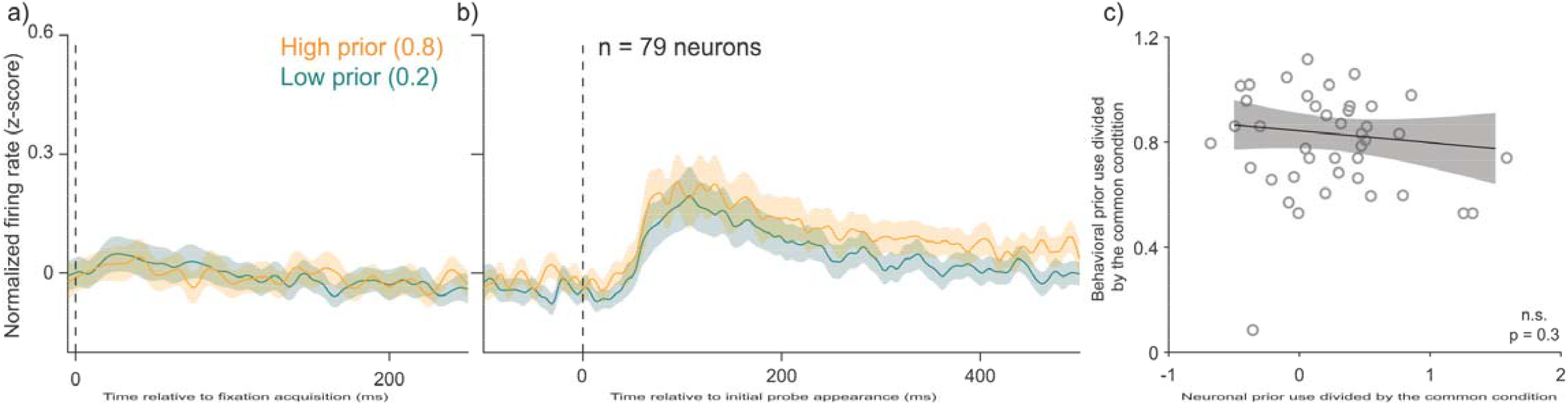
Additional analyses for the no-saccade condition. a-b) Peri-stimulus time histograms in the fixation (a) and probe onset epochs (b) for the no-saccade conditions show overlap in firing rates across the prior conditions. c) There was additionally no correlation between neuronal effect sizes in the probe epoch (0-500 ms from probe appearance) normalized to the common condition and normalized behavioral prior use in the no-saccade, Bayesian comparison condition.

**Figure S8.**
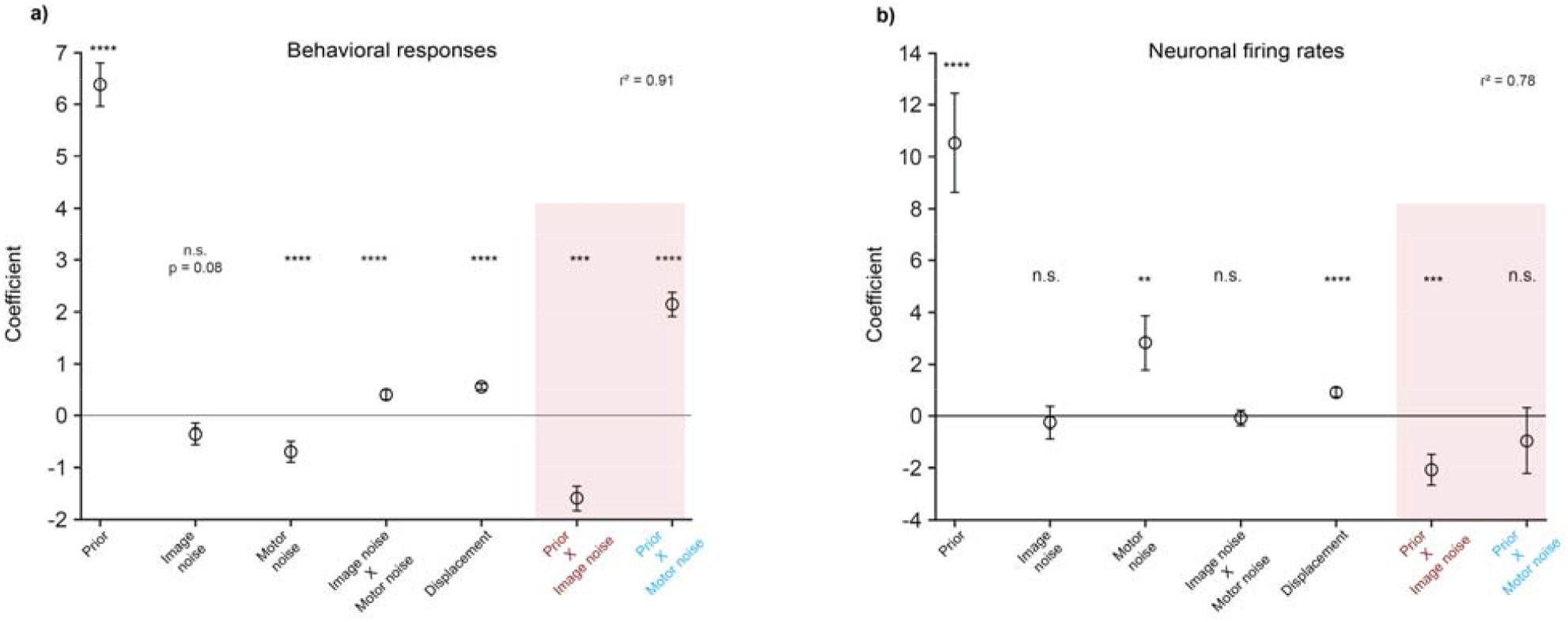
Same as Fig. 5i-j but with all neurons (n = 79) included.

**Figure S9.**
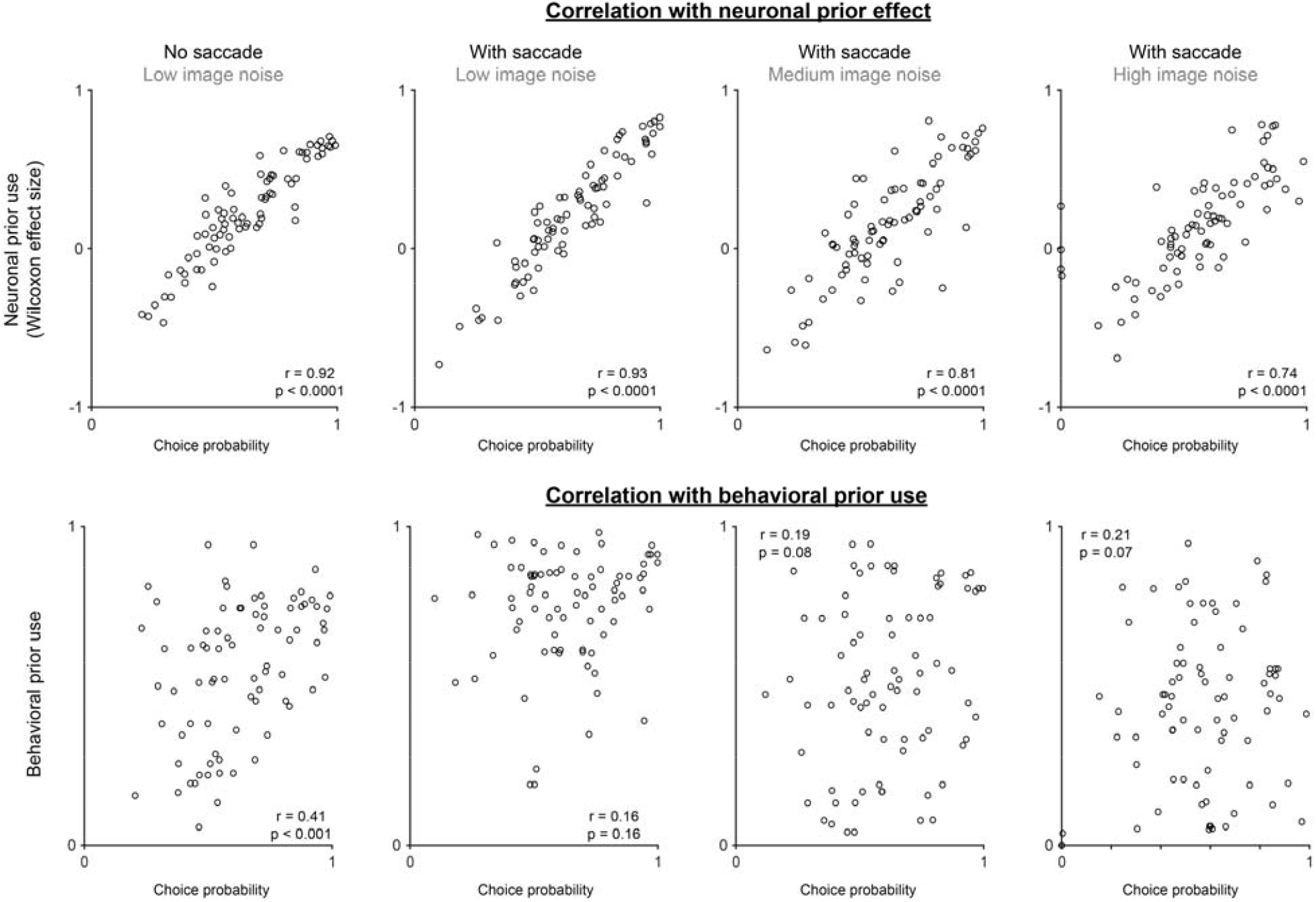
Correlation between choice probabilities for “jumped” vs. “did not jump” responses and prior use. Top row: Correlations with neuronal prior use (Wilcoxon effect size between the high and low prior conditions). Bottom row: Correlations with behavioral prior use (difference in response rates between high and low prior conditions).

